# Early life drought predicts components of adult body size in wild female baboons

**DOI:** 10.1101/2023.01.02.522387

**Authors:** Emily J. Levy, Anna Lee, I. Long’ida Siodi, Emma C. Helmich, Emily M. McLean, Elise J. Malone, Maggie J. Pickard, Riddhi Ranjithkumar, Jenny Tung, Elizabeth A. Archie, Susan C. Alberts

## Abstract

**Objectives:** In many taxa, adverse early-life environments are associated with reduced growth and smaller body size in adulthood. However, in wild primates, we know very little about whether, where, and to what degree trajectories are influenced by early adversity, or which types of early adversity matter most. Here, we use parallel-laser photogrammetry to assess inter-individual predictors of three measures of body size (leg length, forearm length, and shoulder-rump length) in a population of wild female baboons studied since birth.

**Materials and Methods:** Using >2,000 photogrammetric measurements of 127 females, we present a cross-sectional growth curve of wild female baboons (*Papio cynocephalus*) from juvenescence through adulthood. We then test whether females exposed to three important sources of early-life adversity - drought, maternal loss, or a cumulative measure of adversity – were smaller for their age than females who experienced less adversity. Using the ‘animal model’, we also test whether body size is heritable in this study population.

**Results:** Prolonged early-life drought predicted shorter limbs but not shorter torsos (i.e., shoulder-rump lengths). Our other two measures of early-life adversity did not predict any variation in body size. Heritability estimates for body size measures were 36%-58%. Maternal effects accounted for 13%-22% of the variance in leg and forearm length, but no variance in torso length.

**Discussion:** Our results suggest that baboon limbs, but not torsos, grow plastically in response to maternal effects and energetic early-life stress. Our results also reveal considerable heritability for all three body size measures in this study population.

## 1. Introduction

### 1.1 Effects of early-life adversity on body size across species

During development, all organisms must allocate energy toward the body’s many developing systems, including the sensory abilities, the immune system, and growth. When energy is limited, organisms cannot allocate adequate resources to all systems and must develop plastically to survive. Consequently, during early life, adverse environments alter phenotypes by forcing organisms to make developmental trade-offs.

In many taxa, adverse early-life environments, such as famine, drought, or social neglect, have been shown to predict wide-ranging and long-term effects (e.g., blue-footed boobies, *Sula nebouxii*: Ancona & Drummond, 2013; humans, *Homo sapiens*: Brown et al., 2009; humans: Elo & Preston, 1992; humans: Galobardes et al., 2008; humans: Hayward et al., 2013; rhesus macaques, *Macaca mulatta*: Lewis et al., 2000; water pythons, *Liasis fuscus*: Madsen & Shine, 2000; banded mongoose, *Mungos mungo*: Marshall et al., 2017; baboons, *Papio anubis*: Patterson et al., 2021; baboons, *Papio cynocephalus*: Tung et al., 2016). One common effect is slowed growth, which leads to reduced adult body size. For example, in a longitudinal study of the 1958 British birth cohort, children raised in larger families or more crowded homes were shorter than those raised in smaller families or more spacious homes, both in childhood and adulthood (Li et al., 2007). In another study, children raised in Ukrainian orphanages – a source of intense psychosocial stress, but not energetic stress – had stunted growth and low body mass compared to children raised by their families (Dobrova-Krol et al., 2008). Poor early-life environments have also been associated with slower growth and smaller adult body size in wild and captive non-human mammals (wild bighorn sheep, *Ovis canadensis*: Festa-Bianchet et al., 2000; wild roe deer, *Capreolus capreolus*: Pettorelli et al., 2002; captive squirrel monkeys, *Saimiri sciureus boliviensis*: Pucciarelli et al., 2000). For example, wild roe deer born during a time of high population density weighed approximately 15% less as adults than deer born during a time of low population density (Pettorelli et al., 2002).

Whereas growth in humans and ungulates is relatively well studied, analyses of adult body size or growth patterns of most wild mammals are scarce – especially among wild primates. To our knowledge, growth trajectories until or throughout adulthood have only been estimated for wild geladas, baboons, chimpanzees, and mountain gorillas (*Papio cynocephalus*: Altmann & Alberts, 2005; *Gorilla beringei beringei*: Galbany et al., 2017; *Theropithecus gelada*: Lu et al., 2016; *Pan troglodytes schweinfurthii*: Pusey et al., 2005; *Papio anubis*: Strum, 1991). Further, predictors of inter-individual differences in body size or mass have only been assessed for a handful of wild primate populations (Altmann & Alberts, 2005; Berghänel et al., 2015; Johnson, 2003; Pusey et al., 2005; Thompson et al., 2016; Wright et al., 2019, 2020). These studies reveal, for example, that higher social dominance rank was associated with larger body mass-for-age in juvenile and adult female chimpanzees and larger and body size-for-age in adult male gorillas, respectively, but dominance rank did not predict these phenotypes in male chimpanzees or female gorillas (Pusey et al., 2005; Wright et al., 2019, 2020). Among wild baboons, social groups that fed on agricultural crops or human trash gained weight faster and/or reached maximum body mass at younger ages than baboons in naturally-foraging groups (Altmann & Alberts, 2005; Strum, 1991). Furthermore, among naturally-foraging groups of wild anubis baboons, female weight gain slowed during a drought period with low food availability (Strum, 1991). In addition, juvenile Assamese macaques (*Macaca assamensis*) who played more grew less (Berghänel et al., 2015), and semi-free ranging mandrills (*Mandrillus sphinx*) with older or higher-ranking mothers were heavier-for-age than those with younger or lower-ranking mothers (though there was no difference in torso length; Setchell et al., 2001).

Together, this work indicates that primate body size is systematically related to aspects of the social and ecological environments. Furthermore, body size predicts individual fitness in a number of animal species (reviewed in Kingsolver & Pfennig 2004), highlighting the importance of understanding the determinants of adult body size in diverse species. Studies of primate growth in particular can also inform our understanding of human development, health, and disease (Leigh, 2001) and reveal whether the patterns and predictors of growth in non-primate mammals are generalizable to primates, which typically exhibit slower growth rates and slower life histories than other, similarly sized mammals (Charnov & Berrigan, 1993; Pontzer et al., 2014). Finally, a better understanding of the factors affecting primate growth can aid in the effort to combat the urgent population declines and extinction threats facing the majority of primate species (Chapman & Peres, 2021; Estrada et al., 2017).

### 1.2 Effects of early-life adversity on body size in wild baboons

In this study, we use photogrammetry to estimate body size for 127 immature and adult female baboons in the Amboseli ecosystem of southern Kenya. We then test the hypothesis that the energetic and/or psychosocial stress associated with early-life adversity causes small-for-age body size in female baboons. If so, baboons who experienced more early-life adversity will be smaller than those who experienced less adversity. We also provide the first estimate of heritability of body size in a wild primate population.

We test three measures of early life adversity as putative predictors of later-life body size: one that is strongly tied to energetic stress (drought), and two that may be associated with energetic stress, psychosocial stress, or both (maternal loss and cumulative adversity). First, we test early-life drought as a proxy for food availability. Food availability in early life is a well-known predictor of growth, maturation, and body size in primates (Altmann & Alberts, 2005; Altmann et al., 1993; Lee et al., 1986; Leigh, 1994; Mori et al., 1997; Strum, 1991; Sugiyama & Ohsawa, 1982; Whitten & Turner, 2009). Our study population resides in a semi-arid savannah, in which seasonal rainfall patterns are a major predictor of food availability for baboons, who are primarily herbivores (Alberts et al., 2005; Byrne et al., 1993; Post, 1982). Prior studies of this baboon population suggest that baboon growth, fertility, and survival are sensitive to drought and food availability in early life (Altmann et al., 1993; Altmann, 1991; Lea et al., 2015).

Second, we test whether maternal loss (after weaning but prior to adulthood) affects body size later in life. In many mammals, including primates, mothers continue to provide social support, social learning opportunities, and protection against predators after the weaning period (van Noordwijk, 2012). As a result, maternal loss in early life is a strong predictor of shortened adult lifespan in several mammals (e.g., Andres et al., 2013; Foster et al., 2012; Stanton et al., 2020). In the baboon population we studied, offspring whose mothers have died exhibit higher juvenile and adult mortality (Tung et al., 2016; Zipple et al., 2019). Further, offspring whose mothers experienced maternal loss early in life are also less likely to survive to adulthood than offspring whose mothers had not experienced maternal loss (Zipple et al., 2019).

Finally, we investigate the relationship between accumulated early life insults – reflected in a cumulative early-life adversity index that combines six potential sources of adversity – and body size. The six sources include drought and maternal loss, as well as four other aspects of young baboon’s physical and social environment: low maternal dominance rank, which has been associated with smaller size-for-age in prior baboon studies (Altmann & Alberts, 2005; Johnson, 2003); maternal social isolation, which may be associated with psychosocial and/or energetic stress (Silk et al., 2003); large group size, which may increase competition and therefore slow growth; and having a close-in-age younger sibling, which may reduce growth by reducing maternal investment (Tung et al., 2016). Cumulative measures of early-life adversity, such as the Adverse Childhood Experiences index of stressful or traumatic experiences (Felitti et al., 1998), are commonly used in studies of humans because they capture a broad range of factors and provide a continuous and powerful measure of early-life conditions. In baboons, studies measuring a variety of later-life outcomes have found that females who experience greater cumulative early-life adversity exhibit lower adult survival and higher fecal glucocorticoid levels (Patterson et al., 2021; Rosenbaum et al., 2020; Tung et al., 2016). Their offspring are also less likely to survive to adulthood (Zipple et al., 2019). The present study thus lends new insight into a candidate mechanism that links early adversity to fertility and survival later in life: compromised growth and adult body size.

## 2. Methods

### 2.1 Study population and subjects

The study population resides in the Amboseli basin (2°40’S, 37°15’E, 1100m altitude), a semi-arid short-grass savannah ecosystem at the base of Mount Kilimanjaro. In this ecosystem, rainfall is limited and highly variable (Western & Maitumo, 2004). Preferred foods for baboons are most abundant in the rainier months of November-May (Alberts et al., 2005; Altmann, 1998; Post, 1982). During the drier months, from June through October, baboons rely on corms – the underground storage organs of grasses – which require considerable processing effort and thus represent a low-yield fallback food (Alberts et al., 2005; Altmann, 1998, 2009). During droughts, even corms can become limited, sometimes creating extreme food shortages.

The study population consists of wild yellow baboons (*Papio cynocephalus*) with historic and recent admixture with anubis baboons (Papio anubis; Alberts & Altmann, 2001; Vilgalys et al., 2022; Wall et al., 2016). Subjects have been studied since birth by the Amboseli Baboon Research Project. Experienced observers (R. Mututua, S. Sayialel, K. Warutere, L. Siodi) recognize each study subject by sight, and they collect detailed behavioral, demographic, and developmental data from each subject during observation periods several times a week. Female baboons in Amboseli reach sexual maturity in the study population at a median age of 4.5 years and experience their first live birth at a median age of 6.0 years (Onyango et al., 2013). Baboon infants are weaned at approximately 16 months, at which point they are nutritionally independent but still dependent on their mothers for protection, social support, and social learning opportunities (Altmann, 1998). The median lifespan for a female baboon who survives past age 4 is 18.5 years, with a maximum known age of 27.7 years (Tung, Archie et al., 2016).

The study subjects were 127 wild female savannah baboons living in 6 different social groups at the time of data collection (July-December 2019). Subjects were a mix of juvenile, adolescent, and adult females. Birth dates for all subjects were known to within a few days, and subjects ranged in age from 3.3 to 22.6 years, with mean and median ages of 9.9 and 8.7 years, respectively. Of the 127 females in our dataset, 110 had reached sexual maturity, and 89 had given birth to at least one live offspring. Age at menarche and first birth among the study subjects were similar to the population as a whole (4.7 and 6.3 years respectively for the study subjects, as compared to 4.5 and 6.0 years; Onyango et al. 2013).

### 2.2 Measuring early-life adversity

We tested six different early-life environmental variables as predictors of body size: two of them (drought and maternal loss) in separate models, and all six as a cumulative measure in a third model. Each source of adversity is described below.

#### Drought

Guided by prior studies in the Amboseli baboons, we measured both cumulative rainfall and drought (e.g., Beehner et al., 2006; Gesquiere et al., 2008; Tung et al., 2016). We assessed two measures of drought: the proportion of “drought days” during (1) the first year and (2) the first four years of a subject’s life. We define a drought day as a day on which less than 50 mm of rain fell in the previous 30 days (Beehner et al., 2006; Bronikowski & Altmann, 1996; Le Houérou, 1989)(*Supplement: Measuring drought days*). We express these two drought variables as a proportion of drought days – rather than absolute number of drought days – within the first year or first four years of life to accommodate the fact that eight females were slightly younger than four years old at the time of data collection, and therefore did not have as many possible days to experience drought. For these eight females, we calculated the proportion of drought days in the first four years from birth until the mean age at which each female’s photogrammetry data were collected. Because cumulative rainfall and drought overlap in what they measure and produced similar results in our study, we focus on drought in the main text; parallel results for cumulative rainfall are provided in *Supplement: Final dataset and model results*.

#### Maternal loss

Maternal loss was measured as a binary variable indicating whether the subject’s mother died before the subject reached four years of age. Of the eight study subjects who were younger than four years at the time of data collection, one lost her mother before data collection; mothers of the other seven subjects survived past data collection and past their daughters’ fourth birthdays.

#### Cumulative early-life adversity score

The cumulative early-life adversity score was calculated using the six factors identified by Tung, Archie et al (2016) as predictors of adult mortality in female Amboseli baboons. These factors are: (1) experiencing maternal loss before age 4 years; (2) having a close-in-age younger sibling (sibling born <1.5y after subject, approximately the lowest quartile of inter-birth intervals in the population); (3) experiencing high group density at birth (adult group size > 35 at birth, highest quartile for group size in the population); (4) having a low-ranking mother (mother’s ordinal rank > 11 at study subject’s birth, the lowest quartile of ordinal ranks in the population); (5) having a mother with low social connectedness to other females in the first two years of subject’s life (in the lowest quartile of grooming frequency in the population); and (6) being born during a drought year (< 200 mm rain during the first year of life). We used these data to create a cumulative early-life adversity score, which was the total number of adverse events experienced in early life (observed range = 0-4). Because very few individuals had scores greater than three, we grouped all individuals with an adversity score of three or more into a single category. We coded cumulative early-life adversity score as a continuous variable.

### 2.3 Measuring body size via parallel-laser photogrammetry

To measure baboon body size we used parallel-laser photogrammetry, a non-invasive technique that enables repeated measures of body parts, such as the limbs, torso, or tail (Galbany et al., 2016; Lu et al., 2016; Wong & Auger-Méthé, 2018; Wright et al., 2019, 2020). We collected three body size measures: shoulder-rump length (Lu et al., 2016), leg length, and forearm length (Figure 1). All images were taken using an Olympus OM-D EM-5 Mark II mirrorless camera (15.9 megapixel, 3.75μ pixel size) with an M. Zuiko 12-100 mm f/4 IS Pro Digital lens. The lens was fully extended to 100mm during data collection (see *Supplement: Description of housing unit* and *Specifications and calibration of the parallel laser unit*). Our validation of this method demonstrates that the parallel-laser apparatus is highly accurate when compared to manual measurements of inanimate objects (*Supplement: Inanimate object validation*).

**Figure 1.**
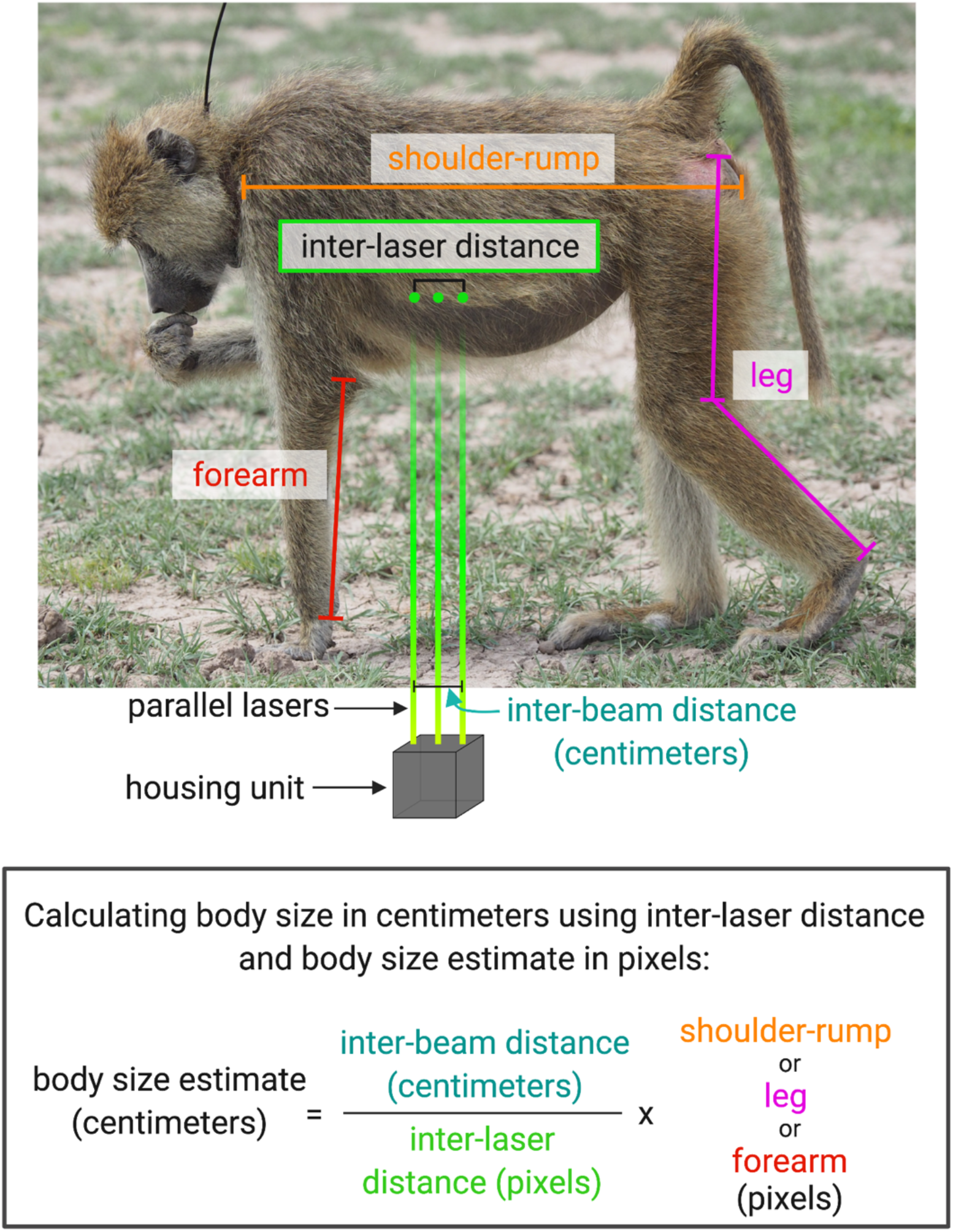
Overview of parallel-laser photogrammetry method. Top: Schematic of image capture and measurement. Parallel laser beams of known inter-beam distance, described in Figure S1, were projected onto the study subject while the subject was photographed. Using ImageJ, we then measured the shoulder-rump length (orange), leg length (pink), forearm length (red), and inter-laser distance (green) in pixels. Bottom: We used this equation to calculate body size in centimeters using the inter-beam distance (in centimeters) between the outer-most laser beams, the inter-laser distance (in pixels) between the two outer-most laser spots projecting onto the study subject, and the body measurement (shoulder-rump, leg, or forearm, in pixels). Adapted from Richardson, Levy et al (2022). Created in BioRender.

Images were taken at a distance of 5-15m from the study subjects, between July and December 2019. To minimize pseudoreplication, measurements of the same subject and same body part were only included if the study subject moved (e.g., took at least one step) between images (see *Supplement: Collecting images*). Images were assessed for inclusion based on the estimated accuracy and precision of a body measurement, given the image characteristics (e.g., focus, distance) and the position of the study subject (e.g., parallax, body part obscured by another baboon; Figure 2; *Supplement: Exclusion criteria)*. Photographers and researchers performing the measurements were blind to the early-life conditions of the study subjects.

**Figure 2.**
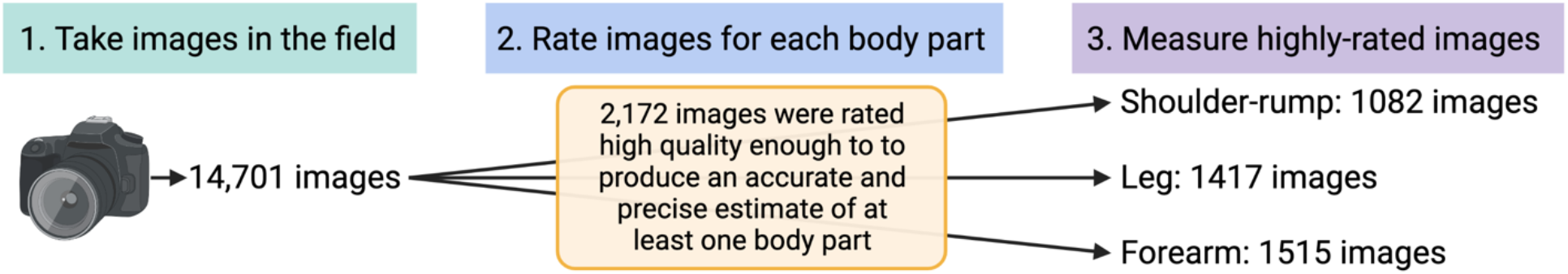
Workflow from image collection (step 1) to image rating (step 2) to image measurement (step 3). For full details of the rating process, see *Supplement*. Created in BioRender.

#### 2.3.3 Measuring images

Landmarks were chosen based on baboon skeletal morphology in an attempt to measure skeletal size alone instead of skeletal size along with muscle and fat (Figure 1; *Supplement: Measuring images* and *Protocol for measuring photogrammetry images*).

To measure the inter-laser distance, we used an automated system that identifies laser spots via brightness and color (Richardson et al., 2022). Of the 2,172 images in our dataset, 1,975 were measured using this automated system. The remaining 197 did not yield inter-laser distance outputs with the automated system so were measured manually in ImageJ. We only measured the distance between the two outer-most laser beams. All hand-measured distances (body measurements and some inter-laser distances) were taken twice and averaged. Observers waited at least one day between making the two measurements.

After collecting the inter-laser distance and body size measurements in pixels, we converted the body size measurements to centimeters. First, we used the inter-laser distance to calculate the number of pixels per 4 cm, and then obtained body size measurements in centimeters by using the following standard equation (see *Supplement: Calibrating the parallel lasers*):

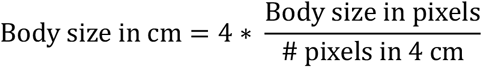

#### 2.3.4 Dataset and variability

Of the 2,172 images used in the analysis, 1,082 yielded shoulder-rump measures for 121 baboons, 1,417 yielded leg measures for 124 baboons, and 1,515 yielded forearm measures for 125 baboons (Table S3, Figure 2; numbers do not sum to 2,172 because some photos yielded measures of multiple body parts). Cumulative early-life adversity score was not available for several study subjects, so the datasets used to test the association between cumulative early-life adversity score and body size were slightly smaller than those for drought and maternal loss (Table S3). Measurement variability was comparable to or lower than those reported in prior studies: within-image percent difference (i.e., difference between two measurements of the same image) was 0.4-1.3%, and within-subject coefficient of variation was 1.9-3.8% (see *Supplement: Image variability and the importance of sample size*).

### 2.4 Using the animal model to assess predictors of body size

Body size is highly heritable in many animal taxa (Hallgrímsson et al., 2002; Mousseau & Roff, 1987; Visscher et al., 2008). To account for genetic effects on body size in our dataset, we ran all our models – both the models that identified the best-fitting cross-sectional growth curve and the models that predicted inter-individual variation in body size – using the quantitative genetic ‘animal model,’ which accounts for relatedness among study subjects. Models were fit in R version 4.1.1 using ASReml-R software version 4, a statistical software package that uses residual maximum likelihood to fit general linear mixed models (Butler et al., 2009).

In all statistical models we included a random effect to account for additive genetic effects, which incorporates the inverse pedigree matrix to represent genetic similarity between individuals in the data set. This approach both allowed us to estimate heritability and controls for the potential confound of relatedness in our estimates of the model fixed effects. Our pedigree included body size for 54 daughters of 31 unique mothers, and 88 (69%) of study subjects had at least one maternal half-or full sister in the dataset. However, the pedigree was incomplete with respect to paternal identity (30 of 127 study subjects had unassigned paternities), limiting our power to fully calculate and control for genetic effects. All our models also included a random effect of maternal identity to estimate combined environmental and genetic maternal effects and to control for the fact that sharing a maternal environment may result in more similar body size above and beyond the similarities arising from relatedness between maternal sisters (Wilson et al., 2010). Finally, to account for the fact that nearly all study subjects were represented by multiple images in the dataset, we included individual identity as a random effect. This random effect estimates repeatability within individuals (Wilson et al., 2010). These three random effects improved the fit of our model (*Supplement: Animal model construction*).

### 2.5 Assessing the shape of cross-sectional growth

To determine the best-fitting cross-sectional growth curve model for shoulder-rump, leg, and forearm lengths of the female baboons in our dataset, we assessed the model fit of three different models of body size as a function of age, using the ASReml-R software. We tested a piecewise linear-linear model with one knot similar to (Pusey et al., 2005), a piecewise quadratic threshold model following (Lu et al 2016), and a quadratic log-log model that approximates logistic growth (see *Supplement: Testing the fit of growth models* for details about these models and how we tested them). Among these three methods, the quadratic log-log model provided the best fit based on R^2^ values (Table S4). As a result, we used the quadratic log-log method to both model a cross-sectional growth curve and to analyze the relationship between early-life adversity and body size. We also used the quadratic log-log method to estimate the ages at which female shoulder-rump, leg, and forearm lengths were at a maximum (i.e., approximating when growth ceases). To do so, we used the coefficients from the model outcomes to calculate the age at which the three models had a slope of 0 (i.e., the apex in the quadratic equations).

### 2.6 Testing environmental and genetic predictors of body size

We fit 18 ‘animal models’ using the quadratic log-log model and the random effects described above to assess whether early-life environments predict adult female body size. Each model had one of three outcome measures: (1) shoulder-rump length, (2) leg length, or (3) forearm length. Each model included one of the four early-life environment predictors: (1) proportion of drought days in the first year of life or (2) first four years of life, (3) maternal loss, or (4) cumulative early-life adversity score (see Supplement for parallel models using cumulative rainfall as a predictor). Twelve models are reported in the main text and all 18 are in the supplement.

Every model included fixed effects of log(age) and log(age)^2^. In the drought and maternal loss models we included an additional predictor of maternal proportional dominance rank at the study subject’s birth because previous studies have shown an association between maternal dominance rank and size-for-age in immature baboons (Altmann & Alberts, 2005; Johnson, 2003). We did not include maternal dominance rank in the three models that contained cumulative early-life adversity score because low maternal rank is also a component of the cumulative score. In the leg length models, we also included a predictor of leg position (slightly bent versus straight), as measurements with the leg slightly bent tended to be shorter than measurements with the leg straight.

Because Amboseli is a hybrid zone between yellow and anubis baboons, we also tested whether level of yellow versus anubis ancestry (i.e., ‘hybrid score’) predicted body size, but the structure of our dataset limited our ability to draw conclusions from this analysis (see *Supplement: Testing whether hybrid score improves model fit*).

### 2.7. Study ethics and data availability

This research was approved by the IACUCs at Duke University and the University of Notre Dame and adhered to all the laws and guidelines of Kenya. Data and are available at the Duke Data Repository (https://doi.org/10.7924/r43r11g2m) and code is available on github (https://github.com/ejlevy/Female-baboon-body-size).

## 3. Results

### 3.1 Cross-sectional growth curves

Based on our quadratic log-log models of cross-sectional growth, the maximum shoulder-rump length was observed in females that were 14.2 years of age, and the maximum leg and forearm lengths were observed in females that were 11.8 and 12.3 years of age, respectively (Figure 3). All three maxima occurred well past the median age at first birth in the population (6 years; Onyango et al., 2013). At their maximum points, our models indicate that average shoulder-rump, leg, and forearm lengths were 47.9, 43.9, and 23.8 cm, respectively.

**Figure 3.**
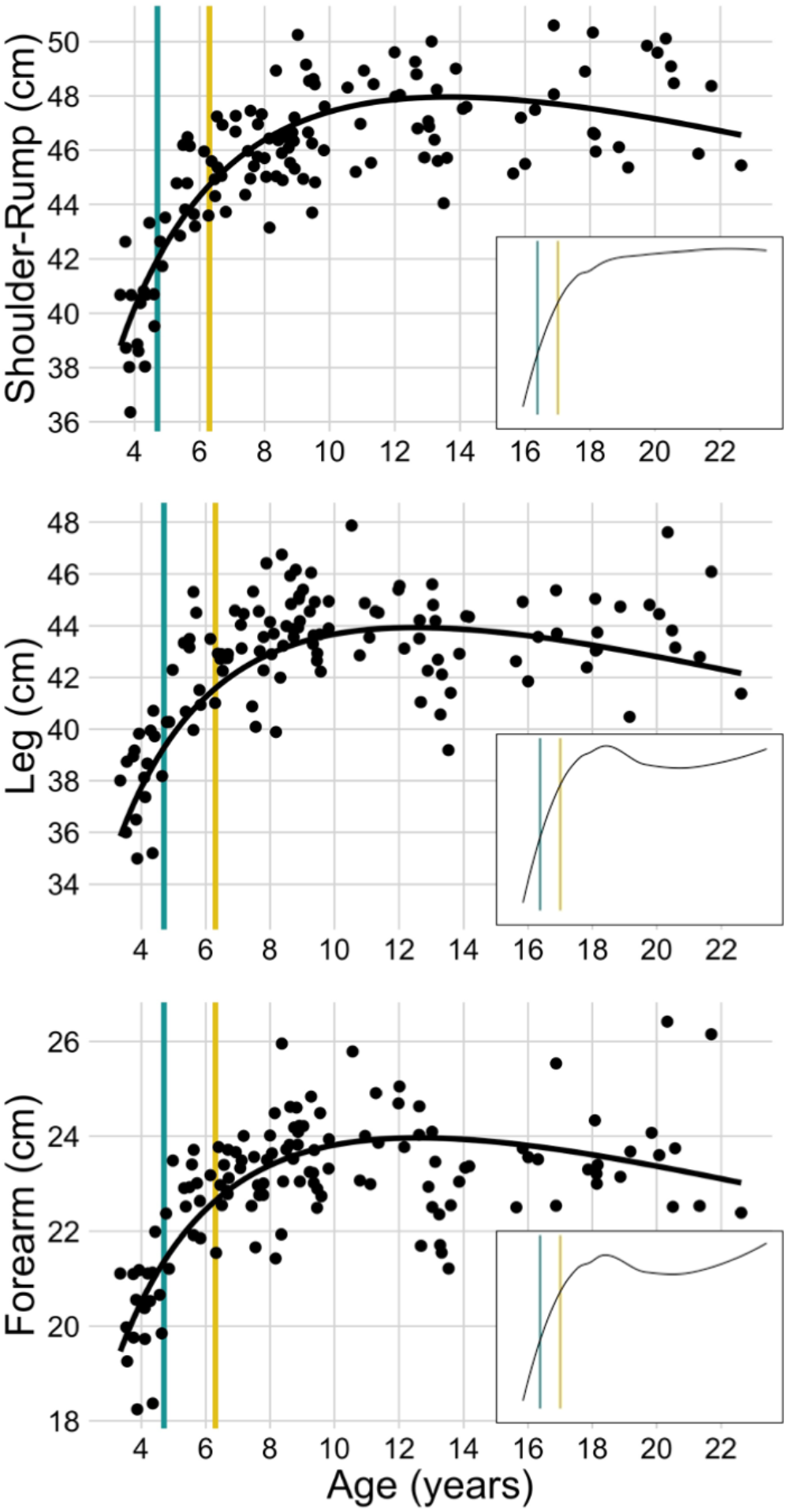
Cross-sectional growth curves for shoulder-rump length (top), leg length (middle), and forearm length (bottom) in female Amboseli baboons. Each point shows mean body size for a unique female, situated along the x-axis at the mean age at which her data were collected. Black lines are the model fit from the quadratic log-log animal models of body size as a function of proportion of drought days in the first four years of life. The vertical lines show median ages at the attainment of two key life-history milestones for females in this data set: menarche (teal; 4.7 years among the study subjects) and first live birth (yellow; 6.3 years among study subjects). Insets show the lowess fits to the same data plotted in the larger figures; for larger plots of the lowess fits, see Figure S5.

Lowess plots for all three body parts show a linear and steep positive slope through 8 years of age, followed by an abrupt change to a weakly positive slope (shoulder-rump) or a mix of weakly positive and weakly negative slopes (leg and forearm; Figure 3 insets, Figure S5). As baboons exhibit determinate growth, we interpret the mix of positive and negative slopes in the lowess plots to be a consequence of small sample size, cohort effects, and/or selection bias for larger females in older age classes.

### 3.2 Early-life adversity and later-life body size: Drought predicts limb length

Both leg and forearm length were predicted by the proportion of drought days in the first four years of life (leg: p = 0.003; forearm: p = 0.005; Table 1). Each additional day of drought in the first 4 years of life predicted leg and forearm lengths that were shorter by 0.015% and 0.017%, respectively. Put another way, a female who experienced drought days in 82.3% of her first four years, which corresponds to 1 SD above the mean of 78.7%, is predicted to have legs and forearms that are 0.83% and 0.90% shorter, respectively, than if she had experienced the mean proportion of drought days. Using our body size maxima, this effect equates to 0.4 cm and 0.2 cm shorter leg and forearm length, respectively. For our study subjects, the proportion of drought days experienced in the first four years of life ranged from 69.1% to 88.2%, indicating a predicted 4.4% (1.9 cm) difference in leg length and 4.8% (1.1 cm) difference in forearm length from the most-to least-affected female. Drought during the first year of life alone had no detectable effect on leg or forearm length (leg: p = 0.37; forearm: p = 0.71; Table S7).

**Table 1.**
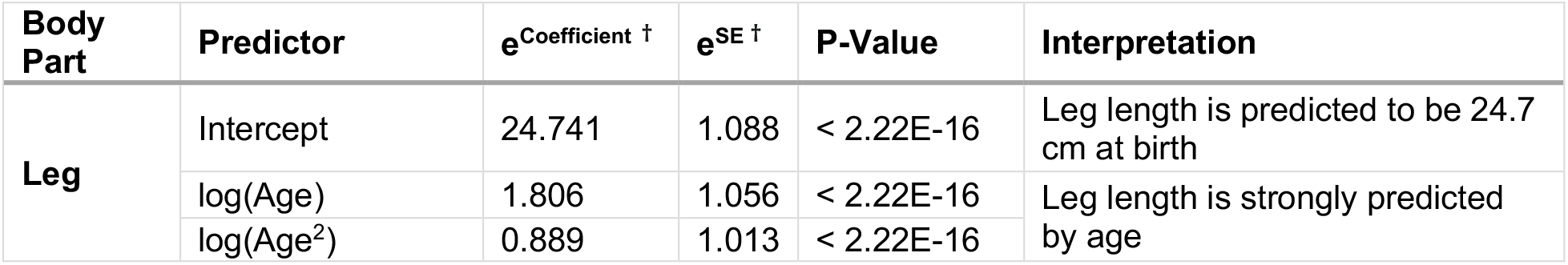
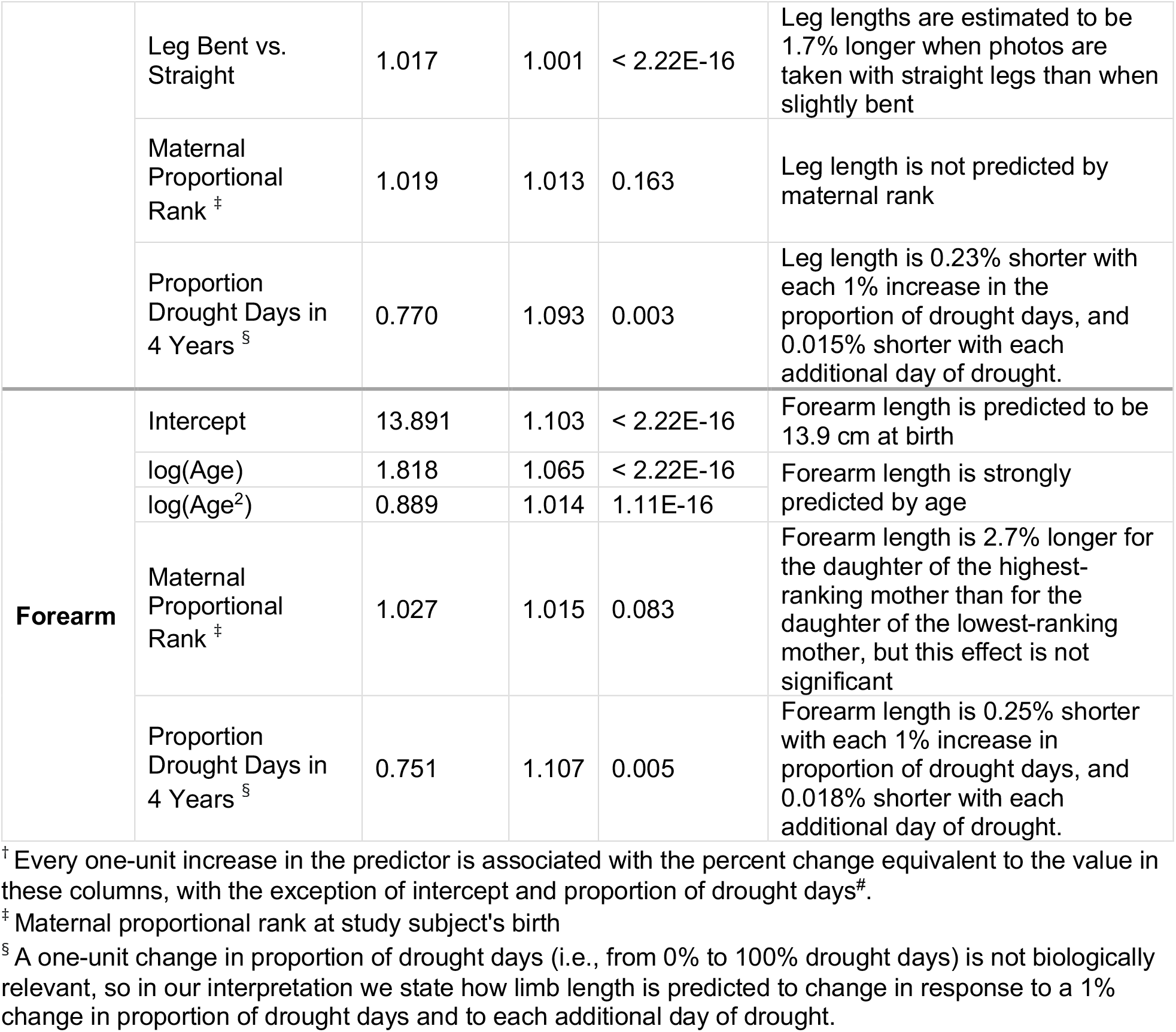
Results of models predicting leg length (top) and forearm length (bottom) as a function of drought in the first 4 years of life (SE = standard error).

Shoulder-rump length was not predicted by drought in the first year or the first four years of life (1^st^ year: p = 0.84; 4 years: p = 0.95; Table S7). Furthermore, none of our body-size measures – shoulder-rump, leg, or forearm length – were predicted by cumulative early-life adversity score or maternal loss (all p > 0.33, Table S7).

### 3.3 Maternal dominance rank and later-life limb length

Forearm length was weakly predicted by maternal rank in the drought and maternal loss models, but the effect did not reach statistical significance. Specifically, the daughter of the highest-ranking mother was predicted to have forearms 2.7-2.9% longer than the daughter of the lowest-ranking mother (p = 0.063–0.095, Table 1, Table S7). Leg length and shoulder-rump length were not predicted by maternal rank (leg: p = 0.11–0.23; shoulder-rump: p > 0.7; Table 1; Table S7).

### 3.4 Body size is highly heritable

Additive genetic variance explained a large proportion of total variance in all three measures of body size in our data set, indicating substantial narrow sense heritability of body size in female baboons. Heritability estimates varied slightly across models and body parts. Heritability of shoulder-rump length was 58%, heritability of leg length was 56-66%, and heritability of forearm length was 36-41% (Table 2, Figure S6, Table S8). Maternal effects also explained a relatively large proportion of variance in limb length, with variance between mothers accounting for 14-22% and 13-17% of variance in leg and forearm length, respectively. In contrast, maternal effects explained almost no variance in shoulder-rump length. The random effect of individual identity, which measures within-individual variation, also explained almost no variance in any body size measures, indicating high repeatability of our photogrammetric measures of body size and therefore relatively small within-individual variation in body size.

**Table 2.** Random effects estimates for the three models that assessed whether the number of drought months a female baboon experienced in her first 4 years of life predicted shoulder-rump, leg, and forearm lengths (ID = identity).

| Body Part | Random Effect | Coefficient | Proportion of Variance | Interpretation |
| --- | --- | --- | --- | --- |
| **Shoulder-rump** | Additive Genetic Effect | 1.27E-03 | 0.58 | Additive genetic variance in the population explains 58% of total variation in shoulder-rump length |
|  | Maternal Effect | 7.56E-10 | 3.44E-07 | Variance between mothers explains < 0.01% of total variation in shoulder-rump length |
|  | ID (Within-Individual Repeatability) | 3.95E-10 | 1.80E-07 | Variance within individuals explains < 0.01% of total variation in shoulder-rump length |
|  | Residual Variance | 9.29E-04 | 0.42 | 42% of variation in shoulder-rump length is unexplained |
| **Leg** | Additive Genetic Effect | 1.11E-03 | 0.56 | Additive genetic variance in the population explains 56% of total variation in leg length |
|  | Maternal Effect | 4.43E-04 | 0.22 | Variance between mothers explains 22% of total variation in leg length |
|  | ID (Within-Individual Repeatability) | 1.81E-06 | 9.16E-04 | Variance within individuals explains 0.092% of total variation in leg length |
|  | Residual Variance | 4.29E-04 | 0.22 | 22% of variation in leg length is unexplained |
| **Forearm** | Additive Genetic Effect | 1.27E-03 | 0.36 | Additive genetic variance in the population explains 36% of total variation in forearm length |
|  | Maternal Effect | 6.22E-04 | 0.17 | Variance between mothers explains 17% of total variation in forearm length |
|  | ID (Within-Individual Repeatability) | 1.71E-10 | 4.77E-08 | Variance within individuals explains < 0.01% of total variation in forearm length |
|  | Residual Variance | 1.69E-03 | 0.47 | 47% of variation in forearm length is unexplained |

Finally, residual variance accounted for 42% of the total variance in shoulder-rump length, 46-47% of the total variance in forearm length, and only 19-22% of the total variance in leg length. The higher residual variances for shoulder-rump and forearm lengths may reflect the fact that the shoulder-rump and forearm measures were noisier than the leg measure (Table S3). See Table 2 and Figure S6 for the random effects estimates from three representative models and Table S8 for the random effects estimates of all 18 models.

## 4 Discussion

### 4.1 Summary of findings

Using cross-sectional body size data from wild female baboons, we found that limb length was stunted by early-life drought, a proxy for low food abundance. Specifically, more exposure to drought in the first 4 years of life was associated with shorter limbs, with a predicted 4 – 5% difference in limb length between the most-affected and least-affected females. contrast, no measure of female body size was predicted by the two measures of early-life adversity that represent combined energetic and psychosocial stressors, cumulative early-life adversity score and maternal loss. Notably, shoulder-rump (i.e., torso) length showed no relationship to any of the early life conditions we examined; only limb length seemed to respond to environmental variation (see ‘Limb growth versus torso growth’ below).

Our findings are best interpreted in the context of the evolution of developmental plasticity. Females may reduce long bone growth in response to early-life energetic stress to allocate limited energy to biological systems that are more crucial for short-term survival and/or long-term fitness. The effects of drought on limb length in this study are of similar magnitude to a study of British children in the 1930s, in which girls whose families were above the median food expenditure had 3.4% longer legs and 2.4% longer trunks than girls whose families were below the median for food expenditure (data extracted from Gunnell et al., 1998 using WebPlotDigitizer). Further, effects of this magnitude may have consequences for health and survival. In a study of over one million Norwegian women born throughout the 1900s, those who were 3% shorter than the height associated with lowest mortality were 1 to 3% more likely to die at any given time, even after controlling for body mass index, age, and cohort effects (Engeland et al., 2003). In nonhuman animals, too, relatively small differences in morphology can be linked to important fitness differences. During an intense drought in the Galapagos Islands, a population of finches (*Geospiza fortis*) experienced a population decline of 85%, with natural selection favoring the largest birds. In this canonical example of natural selection, the differences in body mass and beak size between the pre-drought and post-drought populations were 3.5-5.5% (Boag & Grant, 1981). As Charles Darwin wrote, “What a trifling difference must often determine which shall survive, and which perish!” (Darwin & Wallace, 1858).

Contrary to our expectation, we found that maternal rank did not predict any body size measure except possibly forearm length. In two prior studies of infant and juvenile wild baboons, higher maternal rank predicted larger body mass-for-age (Altmann & Alberts, 2005; Johnson, 2003), a result that aligns with other evidence that higher ranking female baboons have priority of access to food resources (Charpentier et al., 2008; Gesquiere et al., 2018; Post et al., 1980). Our failure to detect an effect of maternal rank on the body size measures used in this study may mean that skeletal growth responds more slowly or subtly to environmental conditions than body mass, which can fluctuate rapidly in response to changing food availability.

Finally, in the first estimates of body size heritability in wild primates, we find that all three measures of body size are highly heritable, in line with estimates of heritability for morphological traits in many other species (Hallgrímsson et al., 2002; Visscher et al., 2008). Further, we found that maternal effects explain variation in limb length but not torso length. The presence of both maternal effects and drought effects for limb lengths, but not for torso length, suggests that limb growth is more plastic in response to early-life environments than growth in the torso.

### 4.2 Limb growth versus torso growth

Our evidence that limb growth (i.e., leg and forearm length), but not torso growth (i.e., shoulder-rump length), is plastic in response to environmental variation agrees with prior literature on humans suggesting that limb length tends to be more affected than torso length in the face of early-life adversity (Billewicz et al., 1983; Bogin et al., 2002; Gunnell et al., 1998; Wadsworth et al., 2002). For example, a longitudinal study of British men born in 1946 found that several early-life factors, such as not being breastfed and having limited energy intake, were associated with shorter adult leg length, whereas fewer of these factors were associated with torso length, and to a smaller degree (Wadsworth et al., 2002). Shorter leg length, but not shorter overall height, has also been associated with lower offspring birth weight and higher incidence of cardiovascular disease and type II diabetes (Chung & Kuzawa, 2014; Gunnell et al., 2003; Lawlor et al., 2004; Smith et al., 2001; Weitzman et al., 2010). During the period when early-life variables are typically measured, humans undergo proportionally more leg growth than torso growth, which may explain greater developmental plasticity in leg length (Gunnell, 2002; Gunnell et al., 1998; Lawlor et al., 2004). Growth plates in the human spine also fuse at later ages than growth plates in the limbs (Albert & Maples, 1995; Buikstra & Ubelaker, 1994; White et al., 2011). If the same timing holds for baboons, there may be more opportunity for compensatory skeletal growth of the torso than of the limbs.

In non-human primates, the relatively few prior studies of skeletal plasticity provide mixed evidence for whether limb length is more plastic than torso length. Japanese macaques (*Macaca fuscata*) living on a small island had shorter limbs relative to their body size than those living on larger islands in Japan, possibly because the population on the small island is near its carrying capacity and has limited food availability (Buck et al., 2018). However, a study of captive capuchin monkeys (*Cebus albifrons*) found no difference between limb and torso growth between individuals raised with and without food restrictions (Fleagle et al., 1975). Much work remains to be done to understand differences between limb growth and torso growth in nonhuman primates.

### 4.3 Heritability and maternal effects

In this study, we estimated additive genetic effects (i.e., narrow-sense heritability) and maternal effects on body size. To our knowledge, this study is the first to do so in a wild primate population. We estimated heritability values of approximately 56-66% for leg length, 36-41% for forearm length, and 58% for shoulder-rump length. These values are within the range reported for the heritability of body size in other non-human animals, which tend to fall between 30 – 70% (Hallgrímsson et al., 2002; Mousseau & Roff, 1987; Visscher et al., 2008). The high heritability of our body size measures suggest segregating genetic variation for these traits and the potential for the population to respond to natural selection.

Maternal effects explained variation in limb lengths (13-22% of the variation) but not shoulder-rump length (Table 2). This difference aligns with our finding of greater plasticity in limb length than in shoulder-rump length in response to drought. In the case of baboons and many other primates, maternal effects can act on energy availability through their effects on learning to forage. Baboons are highly selective omnivores for whom learning to forage requires the presence of tolerant and knowledgeable adults, and mothers commonly fill this role (e.g., Altmann, 1998; Coussi-Korbel & Fragaszy, 1995; King, 1994). While our estimates of heritability and maternal effects are comparable to values seen in other mammals (Maestripieri & Mateo, 2009), we still interpret these values with caution because we were missing paternities in 30 out of 127 study subjects. Further investigation into the source of these maternal effects will be necessary to understand how they may alter the evolutionary potential of these traits.

### 4.4 Growth after sexual maturation

Our data suggest that female baboons may continue to grow for several years past the start of their reproductive careers, though the cross-sectional nature of our data limits our ability to confirm this pattern. Growth past sexual maturity is common in vertebrates: the growth plates of the long bones commonly fuse well after sexual maturity in rodents, ungulates, carnivores, and primates, including humans (Kilborn et al., 2002). For instance, body length in captive chimpanzees and body mass in wild chimpanzees continues to increase for several years past menarche (Hamada & Udono, 2002; Pusey et al., 2005; Walker et al., 2018). Thus, in spite of the limitations of our cross-sectional data, the idea that female baboons continue to grow after sexual maturity is consistent with data from other primates.

If females do continue to grow after they mature sexually, females in energy-limited environments probably face a trade-off between these two energy-intensive processes (Thompson et al., 2016; Whitten & Turner, 2009). This potential tradeoff may explain why the youngest adult females in this baboon population exhibit relatively low fertility, poor infant growth, and reduced infant survival compared to prime-aged females (Altmann & Alberts, 2005; Beehner et al., 2006; Campos et al., 2022; Gesquiere et al., 2018). Relatively poor reproductive performance among young adult females is a common phenomenon in primates (reviewed in (Pusey, 2012)), suggesting that a growth-reproduction tradeoff for females may occur in multiple species.

### 4.5 Limitations and future directions

Several aspects of our study limit the conclusions that we can draw from our findings. First, our data are cross-sectional. While we collected repeated measures of the same female over the span of 1-6 months, this time span is insufficient to measure within-individual growth rates. Instead, we have focused here on body size-for-age in a cross-sectional context.

Collecting longitudinal growth curves for individual baboons will increase our ability to study predictors of inter-individual differences in body size, test for compensatory growth, identify when growth stops, and ask whether female baboons experience a trade-off between growth and reproduction.

Second, although genetic ancestry (i.e., hybrid score) predicts several behavioral and life history phenotypes in this baboon population (Charpentier et al., 2008; Fogel et al., 2021), we were unable to include this predictor in our main models because of limitations on our dataset. Future work expanding the dataset will allow us to learn whether genetic admixture contributes to variation in body size.

Third, we measured three components of skeletal body size, but other components of body size may also be affected by early-life adversity. We did not measure body mass which, in conjunction with body size (i.e., body mass index or body condition) may be more important for female baboon survival and reproduction than body size alone (Gesquiere et al., 2018; Guinet et al., 1998).

And finally, our study is unable to determine whether the magnitude of effects we found are consequential for female baboon foraging efficiency, dominance rank, energy expenditure, and ultimately fitness. Future work will help us resolve how female baboons allocate limited resources to growth and all other developing systems in response to early-life adversity.

## Acknowledgments

Funding for the long-term research is currently through NSF IOS 1456832, NIH R01AG053308, R01AG053330, R01AG071684, R01HD088558, R01AG075914, and P01AG031719; we also thanks Duke University and the University of Notre Dame. In Kenya, our research was approved by the Wildlife Research Training Institute (WRTI), Kenya Wildlife Service (KWS), the National Commission for Science, Technology, and Innovation (NACOSTI), and the National Environment Management Authority (NEMA). We also thank the University of Nairobi, the Institute of Primate Research (IPR), the National Museums of Kenya, the members of the Amboseli-Longido pastoralist communities, the Enduimet Wildlife Management Area, Ker & Downey Safaris, Air Kenya, Safarilink, T. Wango, and V. Oudu for cooperation and assistance. We thank R. Mututua, S. Sayialel, and K. Warutere for assistance in the field; J. Galbany, A. Lu, S. McFarlin, R. Nappi, S. Li, S. Johnsen, S. Solie, J. Harrison, S. Cohen, B. Levy-Cohen, and the Duke Lemur Center for assistance with development and validation of the parallel-laser method; and N. Learn and J. Gordon for database support. For a complete set of acknowledgments of funding sources, logistical assistance, and data collection and management, please visit http://amboselibaboons.nd.edu/acknowledgements/.

