## Supplementary Material for "Early life drought predicts components of adult body size in wild female baboons"

### Measuring drought days

Our definition of a "drought day" follows the drought cutoff suggested by Le Houérou (1989) as a month with rainfall less than twice the mean annual temperature (approximately 50 mm for Amboseli). This statistic has been successfully used by many ecologists to distinguish dry seasons from growing seasons (e.g., Beehner et al., 2006; Bronikowski & Altmann, 1996).

### Specifications and calibration of the parallel laser unit

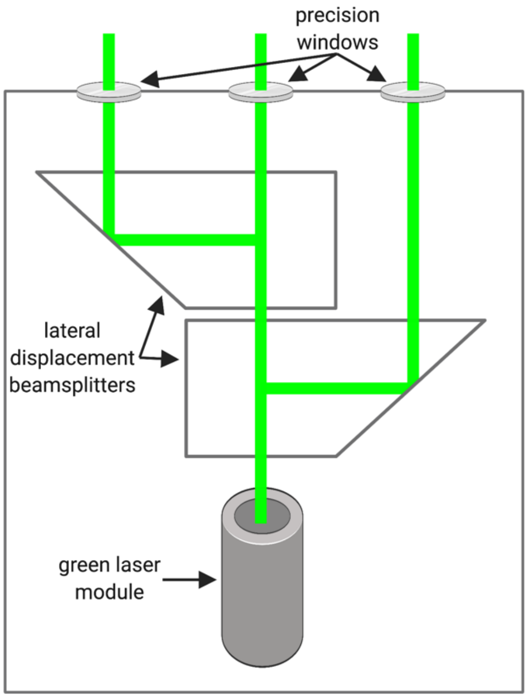

*Figure S1.* Diagram of the parallel-laser housing unit we used to collect body size data. Created in BioRender.

#### *Description of housing unit*

To project parallel laser beams onto the study subjects, we created a housing unit that projects a green laser module (Apinex catalog # AGLM2, <5mW) through two 2 cm lateral displacement beamsplitters (Edmond Optics catalog # 47189; Figure S1). The two beamsplitters were stacked in the housing unit such that the beam from the laser module was split into 3 beams, each approximately 2 cm apart, with the farthest two beams approximately 4 cm apart (Figure S1). This equipment was housed in an aluminum case with plastic front and back. The three laser beams exited the case through 3 mm-thick precision windows (Thorlabs, catalog # WG10530-A). The laser module was powered by two AA batteries mounted to the outside of the case.

*Calibrating the parallel lasers*

The two beamsplitters produced nearly parallel beams, but we were able to measure slight yet predictable deviation from parallel. Consequently, when the study subject was farther from the camera, the physical distance between the two outer-most laser spots was slightly larger than when the study subject was closer to the camera. To assess and correct for this deviation, we created an equation that converted the inter-laser distance (i.e., the number of pixels between the two farthest laser spots) into the number of pixels, at that distance, that would represent a true 4 cm distance. To do this, we took images of the laser beams projecting onto a flat surface that was marked with a true 4 cm distance. In those images we digitally measured (1) the number of pixels between the two outer-most beams as well as (2) the number of pixels spanning the true 4 cm distance. We collected these two measurements repeatedly from images taken 2-20 m from the flat surface. Changing our distance to the flat surface allowed us to determine the relationship between the distance from the camera to the flat surface and the distance between the laser spots on the surface. Using these data, we converted each image's inter-laser distance in pixels to 4 cm in pixels using the following equation: 4 cm in pixels = -0.99 + 0.99*(inter-laser distance in pixels). We then converted body length measurements from pixels to centimeters (Figure 1 in main text).

### Inanimate object validation

To assess the accuracy of the parallel-laser method, we carried out a validation by comparing measurements of inanimate objects obtained via parallel-laser photogrammetry to measurements of those same objects obtained by hand using a measuring tape. We photographed inanimate objects with defined endpoints to increase the accuracy and precision of the hand measurement, so that the hand measurement approaches the 'true' measurement. Images of several objects (garage door window, fence posts, piece of paper) were taken from 5, 10, and 15 meters. We measured 1 image from each distance and performed 1-3 measurements on each image. Inter-laser distance was obtained via the automated method (see Measuring images below), except for 1 image (piece of paper from 15 m) for which the automated method did not work and was therefore measured manually. We calculated the <u>absolute difference</u> between manual and parallel-laser measurements. We also calculated the <u>percent difference</u> by dividing the absolute difference by the manual measurement and multiplying by 100.

Results from this validation demonstrate that the parallel-laser apparatus is highly accurate (Table S1). Parallel-laser measurements tended to be within 0.2 cm of the manual measurements (mean = -0.1 cm; SD = 0.2 cm). This level of error resulted in a percent difference that ranged from -2.7% to 0.4% (mean = -0.6%; SD = 0.8%) and depended on the size of the object measured (i.e., i.e., the percent difference was larger for smaller objects than for larger objects). Measurements taken via parallel-laser photogrammetry have similar absolute differences from manual measurements, regardless of the size of the object being measured, and absolute differences between manual and parallel-laser measurements were highly consistent across 5, 10, and 15 m distances (Table S1). However, the parallel-laser measurements were consistently – although only slightly – larger than the manual measurements, indicating a slight systematic bias towards larger size measurements with the parallel-laser method.

**Table S1.** Results of validation using inanimate objects, averaged across distance from parallel-laser apparatus to object (5, 10, 15m).

| Object | Measurement | Mean manual (cm) | Mean parallel-laser (cm) | Difference[a] | % Difference[b] | Parallel-laser SD[c] |
|---|---|---|---|---|---|---|
| Door | Width of post between windows | 8.6 | 8.83 | -0.2 | -2.7 | 0.0 |
| | Width of inside frame of one window panel[d] | 34.3 | 34.48 | -0.2 | -0.5 | 0.1 |
| | Width of outside frame of one window panel | 36.4 | 36.42 | 0.0 | -0.1 | 0.2 |
| | Width of two window panels[4] | 78.2 | 78.30 | -0.1 | -0.1 | 0.2 |
| Fence | Width of one plank | 13.6 | 13.68 | -0.1 | -0.6 | 0.0 |
| | Width of two planks | 28.1 | 28.20 | -0.1 | -0.4 | 0.1 |
| | Width of four planks | 56.7 | 56.84 | -0.1 | -0.3 | 0.2 |
| Paper | Height | 21.6 | 21.62 | 0.0 | -0.1 | 0.0 |
| | Width | 27.9 | 27.90 | 0.0 | 0.0 | 0.1 |
| **Mean of means** | | | | **-0.1** | **-0.5** | **0.1** |

[a] Difference calculated as Manual measurement – Parallel laser measurement
[b] % Difference calculated as Difference/Manual*100
[c] SD of the three parallel-laser measurements collected for that particular object.
[d] Manual measurements for this object were challenging, and may be less accurate than other measurements for other objects.

### Collecting images

Nearly all images were taken by co-author Anna Lee; those not taken by AL were taken by first author Emily J. Levy. Most images were taken of subjects whose identities were known by the photographer at the time the image was taken. In cases in which the identity of the focal baboon was not known while photographing her, identities were confirmed based on visual inspection of the images soon after by experienced observers (either co-author I. Long'ida Siodi, or J.K. Warutere or R.S. Mututua). Images in which the identity of the study subject was uncertain were excluded from the dataset.

### Exclusion criteria for images

Exclusion criteria depended on the body part being measured (Table S2).

Images that were not excluded from shoulder-rump measurements were given shoulder-rump ratings of 3.5, 4, 4.5, or 5, with higher ratings reflecting better image quality and body position. We also assessed whether a more exclusive dataset – of shoulder-rump ratings of 4 and higher – reduced error in the analysis. With the larger more inclusive dataset, we detected more variance, but a lower proportion of that variance was residual variance compared to the more exclusive dataset, so we report results based on the more inclusive dataset in our analysis.

Images that were not excluded from leg measurements were given leg ratings of 1 or 2, with 1 indicating that the leg was slightly bent, and 2 indicating that the leg was straight.

All images that were not excluded from forearm measurements were measured; there was no rating system for the forearm measurement.

**Table S2.** Guidelines for excluding photogrammetry images; occurrence of one or more exclusion criteria in an image precludes its measurement for that particular body part.

| **Exclusion criterion** | **If answer is yes, exclude measurement of these body parts (Exceptions in parentheses)** [a] |
|---|---|
| The subject is blurry. Slight blurriness is ok if you feel confident in your ability to identify the landmarks and lasers. | SR[b], Leg, Forearm |
| The lasers points are impossible to identify. Before excluding the image, try adjusting the color balance in the Lightroom app. If you still can't see the lasers, you can't measure it. | SR, Leg, Forearm |
| Parallax error; the subject's spine is not perpendicular to the camera. | SR, Leg, Forearm (but can include Leg and Forearm if parallax is very slight) |
| The subject is walking | SR |
| The subject's limbs are not perpendicular to ground and spine. This is especially important for the limbs facing the camera but can also be affected by torso twisting due to the other two limbs being angled. If only slightly off perpendicular, ok to include. | SR, Leg |
| Leg facing the camera is bent more than 45 degrees at the knee and/or heel is lifted off the ground, indicating a flexed ankle. | SR, Leg (but can include SR if femur is perpendicular to spine) |
| You can see both ischial callosities or the far side of the paracallosal skin. You can include if you can see a tiny bit of the far ischial callosity *above* the closer callosity – this happens sometimes if the subject is relatively close to you, because the camera is angled down onto the subject. Note: if the female has a large sexual swelling, it may be difficult to apply this criterion. If so, photo should be excluded. | SR, Leg (but can include Leg if the back callosity is barely visible) |
| Arm facing the camera is bent (e.g., subject is putting food in her mouth). | SR (but can include SR if shoulder is in proper position) |

| | |
|---|---|
| The subject has a noticeably curved or hunched spine (sometimes seen when baboon is looking very far down). If the subject's spine always looks like that (e.g., because of age), then include. | SR |
| Twisted torso: The subject's head is rotated to look laterally, such that you can see both eyes (if turned toward you) or no eyes (if turned away from you). Looking laterally can cause the torso to twist. You can also identify twisted torsos if you see both callosities and both shoulders. | SR |

[a] Parentheticals indicate instances in which the exclusion criterion is slightly different for a particular body part.
[b] SR = shoulder-rump

### Measuring images

Images of sufficiently high quality were measured manually by EJL using ImageJ (Schneider et al., 2012). By having only one person measure images, we eliminated between-observer error, which tends to be larger than within-observer error (Barrickman et al., 2015; Berghänel et al., 2015; Breuer et al., 2007; Galbany et al., 2016; Lu et al., 2016; Wright et al., 2019).

For shoulder-rump measurements, images were first rotated so that the study subject's spine was parallel to the top of the computer screen. A rectangle was then drawn around the subject, with one side extending to the outer-most point of the ischial callosity and the other side to the bony point of the shoulder, slightly inward from the edge of the fur. The width of this rectangle was used as the shoulder-rump length (orange line in Figure 1 in main text; see also Figure S7). Leg measurements were taken by first drawing a line from the outer-most curve of the heel to the inside of the knee. Using the exact same location for the knee landmark, a second line was drawn from there up to the top-most point of the ischial callosity (pink line in Figure 1 in main text; see also Figure S8). These two lengths were then added together. Forearm measurements were taken by drawing a line from the elbow to the indent just below the protrusion at the wrist marking the end of the ulna (red line in Figure 1 in main text, see also Figure S10). All measurements for all body parts were taken twice and averaged. We increased the independence of measurements by spacing repeated measurements by at least one day.

### Image variability and the importance of sample size

Several steps in the parallel-laser photogrammetry process can introduce variability into body size measurements (Galbany et al., 2016; Richardson et al., 2022). Common sources of variability are parallax, body position, and lack of repeatability when identifying landmarks, both in the same image and across images. We assessed two types of variability in our datasets: (1) <u>within-image difference and percent difference</u>, which capture the repeatability of the observer measuring the same image twice, and (2) <u>within-subject standard error</u> (SE) and <u>within-subject percent coefficient of variation</u> (CV), which capture the repeatability of measurements across images of the same study subject (Table S3).

Within-image percent difference was similar to prior studies that used parallel-laser photogrammetry to measure body size in wild primates, which report within-image percent differences of 0.5% - 2.0% (Breuer et al., 2007; Galbany et al., 2016; Lu et al., 2016; Wright et al., 2019). Within-subject SE was twice as high for the shoulder-rump dataset as for the leg and forearm datasets, indicating that the shoulder-rump measure was slightly less repeatable across images of the same baboon. Within-subject percent CV values were comparable to variability in prior studies, which report within-subject percent CVs of 1.0% - 5.0% (Galbany et al., 2016; Lu et al., 2016; Wright et al., 2020). The within-image percent difference and within-subject percent CV were likely larger for the forearm measurement because it is approximately half as long as the leg and shoulder-rump measurements.

**Table S3.** Descriptive information about the body size datasets used in analysis.

| Body part | # subjects[a] | # images | Mean # images/subject | Age span (years)[b] | Within-image difference (cm)[c] | Within-image % difference[d] | Within-subject SE | Within-subject % CV[e] |
|---|---|---|---|---|---|---|---|---|
| Dataset used to assess shape of cross-sectional growth curve | | | | | | | | |
| Shoulder-rump | 121 | 1082 | 8.9 | 0.16 | 0.20 | 0.44 | 0.50 | 2.97 |
| Leg | 124 | 1417 | 11.4 | 0.17 | 0.17 | 0.40 | 0.28 | 1.99 |
| Forearm | 125 | 1515 | 12.1 | 0.17 | 0.30 | 1.32 | 0.27 | 3.77 |
| Dataset used to assess effects of rainfall, drought, and maternal loss | | | | | | | | |
| Shoulder-rump | 121 | 1082 | 8.9 | 0.16 | 0.20 | 0.44 | 0.50 | 2.97 |
| Leg | 123 | 1416 | 11.5 | 0.17 | 0.17 | 0.40 | 0.28 | 1.99 |
| Forearm | 124 | 1513 | 12.2 | 0.18 | 0.30 | 1.32 | 0.27 | 3.79 |
| Dataset used to assess effects of ELA score | | | | | | | | |
| Shoulder-rump | 117 | 1055 | 9.0 | 0.16 | 0.19 | 0.43 | 0.49 | 2.96 |
| Leg | 119 | 1389 | 11.7 | 0.17 | 0.17 | 0.40 | 0.27 | 1.94 |
| Forearm | 120 | 1475 | 12.3 | 0.17 | 0.31 | 1.33 | 0.27 | 3.79 |

[a] Early-life data were missing for several study subjects, so the dataset for the cross-sectional growth curve analysis (top) is slightly larger than the dataset for the rainfall, drought, and maternal loss analyses (middle) and the cumulative ELA score analysis (bottom).
[b] Age span: Each study subject's images were taken over a time span of 0-0.5 years; here, we calculate the mean of that time span across study subjects, in years.
[c] Within-image difference: First, we calculate the difference between measurement 1 and measurement 2 of the same image and designate this as the within-image difference for a particular image. We then take the mean of within-image differences across all images in the dataset.
[d] Within-image % difference: First, we calculate the difference between measurement 1 and measurement 2 of the same image and designate this as the within-image difference for a particular image; then we divide the within-image difference by the mean of the image's 2 measurements, and multiply by 100 to calculate the image's within-image percent difference. We then take the mean of within-image percent difference across all images in the dataset.
[e] Within-subject % CV: First we calculate the SD across all photos of the same study subject divided by the mean of all photos of that same subject, and multiply by 100 to express as a percentage. We then take the mean of those percentages across study subjects. Note that images were taken over a time span of 0-0.5 years.

In any normally distributed population, larger samples of that population will have smaller variation (i.e., standard error) than smaller samples. Likewise, larger samples will better approximate the true mean of the population. In a similar vein, more photogrammetry images of a particular baboon should both reduce within-individual standard error and better approximate that individual's "true" body size.

To learn whether and to what extent our within-individual body size estimates improved as number of images increased, we performed a random sampling analysis of all study subjects for whom we had at least 15 images for a particular body measurement. We had 14 females for whom we had at least 15 images with shoulder-rump measurements, 35 females for whom we had at least 15 images with leg measurements, and 49 females for whom we had at least 15 images with forearm measurements). Using these data, for each individual baboon we then randomly sampled measurements from 15 images without replacement; then 14 images from the original image collection, without replacement; then 13 from the original image collection, without replacement, and so on. For example, when sampling 15 images out of a possible total of 23, each of those 15 images was unique; but when a separate sample of 14 images was drawn, we again sampled from the original full set of 23 images, meaning some images could re-occur in the 15 sample subset and the 14 sample subset. From these samples of images, we then calculated the mean and percent CV for each individual baboon's measurements. The resulting dataset had, for each female, 14 different means (one from the random sampling of 15, one from the random sampling of 14, and so on) and 14 percent CVs (mean and CV cannot be calculated from a sample of 1).

We used this dataset to visualize two relationships. First, we asked how the number of images we use to estimate the size of a body part affects the variation in our estimate. We found that across all individuals, our measure of variation (percent CV) does not decline when we had more images of a particular individual, as evidenced by the fact that the mean CV does not decline as number of images increases (Figure S2, top 3 panels). In fact, for the shoulder-rump and forearm measurements, at very small numbers of images (2-5), percent CVs were lower than at larger numbers of images. At these small number of images, the percent CVs were also more variable, as evidenced by the larger ranges in these data (i.e., taller violins in plots) for smaller numbers for the shoulder-rump and forearm measurements (Figure S2, top left and top right panels). We interpret this as an indication that larger numbers of images for each study subject do not reliably reduce within-subject variability.

Second, we asked how the number of images we use to estimate the size of a body part affects our ability to estimate a baboon 's "true" body size. Although we cannot know a baboon's true body size, we expect that the more measurements we have of an individual, the closer we are to her true size. To assess this, we calculated each baboon's mean measurement for each sample size of images, and then for each sample size we compared the estimate of her mean at that sample size compared to her mean at a sample size of 15; we expressed this comparison as a percentage difference. As we expected, the larger the samples size of images we used to estimate a body size measurement, the lower the variation around the estimate of the mean body size (Figure S2, bottom panels), suggesting that larger numbers of photos allowed us to more accurately measure a female's true body size.

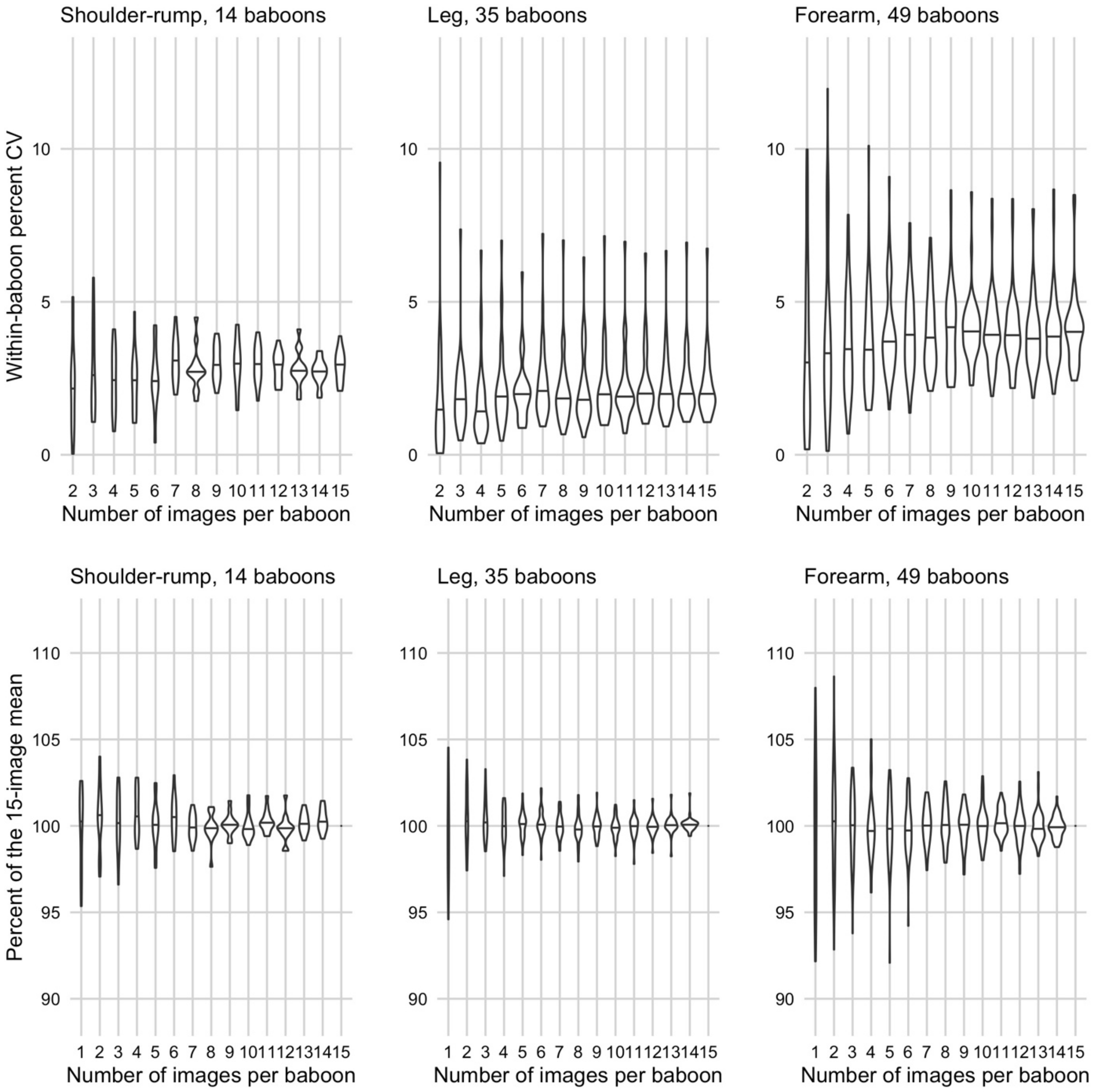

**Figure S2.** Top panels: Within-baboon percent CV as a function of the number of images used to estimate the size of the body part, for shoulder-rump (left), leg (center) and forearm measures (right). Across all study subjects, percent CV does not decline with more images, but it does become less variable across study subjects for shoulder-rump (left) and forearm (right) measurements, indicating that larger sample sizes for each study subject does not reliably decrease within-baboon variability. Bottom panels: Within-baboon mean measurements as a function of the number of images used to estimate the size of the body part, for shoulder-rump (left), leg (center) and forearm measures (right). The y-axis is a baboon's mean measurement for a particular number of images divided by her measurement with a sample size of 15, then multiplied by 100. A value of 100 on the y-axis means that her mean measurement at that particular number of images is identical to her mean measurement with 15 images. As sample size increases, body size estimates more closely approximate the 15-image mean, as evidenced by less variability in the dataset as the number of images per baboon increases.

### Testing the fit of growth models

To assess how to model cross-sectional growth, we tested our body size data with three types of models: a piece-wise linear model, a quadratic threshold model, and a quadratic log-log model. All models were run in R version 4.1.1 as linear mixed-effects animal models using the ASReml-R software version 4 (Butler et al., 2009). The animal model allows us to control for possibility of related individuals having similar body sizes due to shared genotypes; these models also estimate heritability. Each model included a random effect of individual baboon, because the unit of analysis was a single image's measurements and we had repeated measures for most study subjects. Each model also included random effects of maternal identity and breeding value, which was estimated in part based on an inverse pedigree matrix that we supplied.

First, we tested a piecewise linear-linear model with one knot, which assumes that animals grow linearly until they abruptly stop growing; a version of this model, with two knots instead of one, was used to assess growth in wild chimpanzees (Pusey et al., 2005). Second, we tested a quadratic threshold model that assumes a quadratic growth trajectory before the knot and has a slope of 0 after the knot. This quadratic threshold model has been used to assess growth curves for gelada monkeys (Lu et al 2016). The quadratic threshold model approximates Gompertz and von Bertalanffy logistic growth models, which represent the traditional approaches to modeling growth, but the quadratic threshold model is more amenable to a mixed-effects modeling approach than the Gompertz and von Bertalanffy logistic models. Finally, we tested a quadratic log-log model. We reasoned that log-transforming both age and body size would be the best approach to detecting logistic growth, the growth process that is assumed by both the Gompertz and von Bertalanffy growth equations. In addition, the quadratic log-log model allows for intuitive interpretation of model results.

*Creating the models*

For all three growth models, the outcome variable was body size, the fixed effect was age, and the random effect was individual identity. To test these three models against each other, we first optimized the knot location for the two piecewise models (piecewise linear-linear and quadratic threshold). To do this optimization, we ran each model iteratively, each time changing the knot location by 0.1 years from the minimum to the maximum age range of the study subjects in the dataset. We extracted the AIC score for each of these models and chose the knot location that yielded the model with the lowest AIC score. This model-optimizing process was repeated three times, once for each body measurement (shoulder-rump, leg, forearm). For example, to assess the optimal knot location for the piecewise linear-linear model in predicting forearm length, we ran the piecewise linear-linear model with the forearm dataset – first with the knot at 3.3 years, then at 3.4 years, then 3.5, etc. We then found that the AIC value was lowest for the model in which the knot location was at 5.5 years. The fit of this model (i.e., $R^2$ values in Table S4) was then compared to the model fit of the quadratic threshold model (for which the optimal knot location was assessed using the same method) and the quadratic log-log model.

For the quadratic log-log model, which was not piecewise and thus did not require this optimization process, we included log(age) and log($age^2$) predictors. For all three types of model (piecewise linear, quadratic threshold, and quadratic log-log), the models of leg length also included a leg image rating as a predictor, as ratings with a score of 1 (leg slightly bent) tended to be yield shorter leg measurements than ratings with a score of 2 (leg straight) (see *Exclusion* *criteria* above). We also tested whether including image rating for shoulder-rump (see *Exclusion criteria* above) in our model as a predictor variable improved models of shoulder-rump length, but it did not (ΔAIC = 12.0), probably because lower ratings represent more noise in the measurement as opposed to a bias toward longer or shorter lengths.

*Testing the growth models: optimal knot locations, visualization, and $R^2$*

For the shoulder-rump measurement, the optimal knot locations were 6.4 years for piecewise linear-linear and 7.9 years for quadratic threshold. For the leg measurement, the optimal knot locations were 5.8 years for piecewise linear-linear and 5.0 years for quadratic threshold. For the forearm measurement, the optimal knot locations were 5.5 years for piecewise linear-linear and 4.7 years for quadratic threshold. We suspect that these models, especially those of leg and forearm length, provide relatively poor fits to the data. First, for the quadratic threshold models of the leg and forearm datasets, the quadratic component produced a 'U' shaped curve between ages 4 and 5 years (Figure S3, middle-right and bottom-right panels; 'U' inflection point at approximately 4 years). Second, the knot locations can be thought of as estimates of the age at which growth ceases. However, visual inspection of the raw data indicates that females between 7 and 10 years of age are consistently larger than 5-6 year-olds, suggesting that growth continues after the optimal knot locations. While we cannot rule out a strong cohort effect or genetic effect on body size as explanations for the consistently larger size of 7-10 year-olds, these explanations seem unlikely given the magnitude of the size difference between these age classes. See Figure S3 for visualizations of these optimized piece-wise models.

We then calculated $R^2$ for each model output to assess how well each type of model explained variation in body size. For all body size measures, $R^2$ values were highest for the quadratic log-log models, so we used the quadratic log-log models in all further analyses (Table S4).

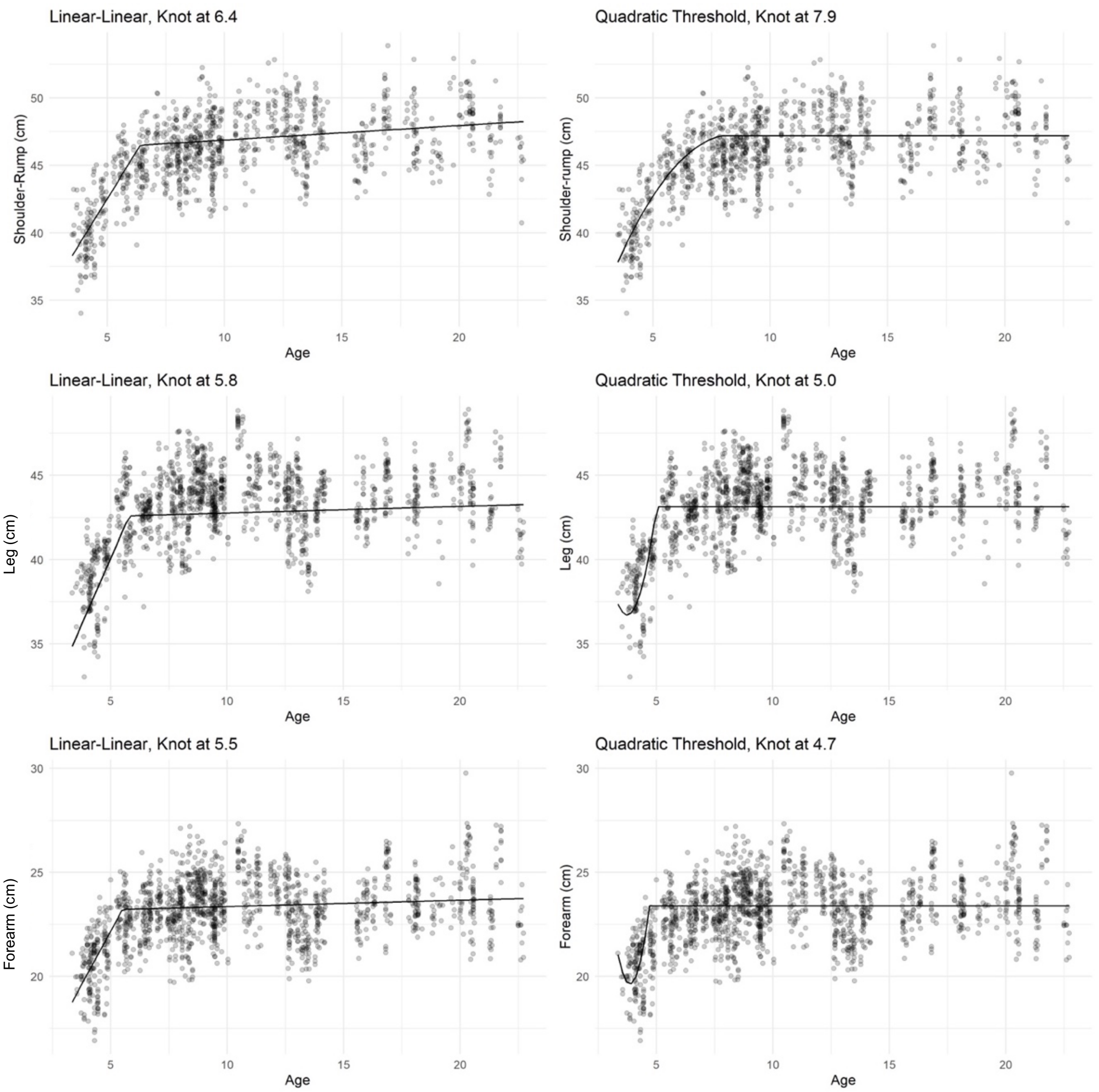

**Figure S3.** Visualizations of the model results from the piecewise linear piecewise models (left side) and quadratic threshold models (right side) for shoulder-rump (top row), leg (middle row), and forearm length (bottom row). Each datapoint represents a single image, and the black lines show the model predictions.

**Table S4.** Identifying the best model of cross-sectional growth curves in female baboons; values are the $R^2$ outputs for models of each body part (shoulder-rump, leg, forearm) modeled in three different ways.

| | | Piecewise linear-linear | Piecewise quadratic threshold | Quadratic log-log |
|---|---|---|---|---|
| Marginal $R^{2\ a}$ | Shoulder-rump | 0.429 | 0.522 | 0.548 |
| | Leg | 0.232 | 0.420 | 0.454 |
| | Forearm | 0.307 | 0.285 | 0.328 |
| Conditional $R^{2\ b}$ | Shoulder-rump | 0.781 | 0.783 | 0.795 |
| | Leg | 0.853 | 0.871 | 0.879 |
| | Forearm | 0.660 | 0.669 | 0.680 |

[a] Marginal $R^2$ represents variation explained by the fixed effects only (age for shoulder-rump and forearm models, age and leg rating for leg models)
[b] Conditional $R^2$ represents variation explained by fixed effects (age for shoulder-rump and forearm models, age and leg rating for leg models) and random effects.

### Assessing inter-individual variation in body size using the animal model

To assess model fit with and without relatedness information, we chose one representative model (leg length as a function of drought days in the first four years of life) and ran three alternative models that either (1) did not include the pedigree, (2) did not include the maternal identity, or (3) did not include both pedigree and maternal identity. The original model, which included both pedigree and maternal identity, was preferred over all of these alternative models ($\Delta$AIC = 3.3-19.1).

ASReml-R iteratively maximizes likelihood in a stepwise fashion, and we ran 400 iterations of each model. For two of our 18 models of body size (those predicting leg length as a function of drought days in the first year of life and leg length as a function of maternal loss), the model output warned of a convergence error. In these two cases, increasing the number of iterations to 1,000,000 did not change model outputs or the warning, so we report the model outputs with 400 iterations here.

Because two common measures of dominance rank – ordinal and proportional – differ in their ability to predict phenotypes, we assessed model fit with both measures of maternal rank using one representative model (leg length as a function of drought days in the first four years of life)(Levy et al., 2020). The model with maternal proportional rank was preferred and was thus used in all of our other analyses ($\Delta$AIC = 6.6).

### Testing whether hybrid score improves model fit

The Amboseli baboon population is a majority-yellow baboon population that includes descendants of historical admixture prior to the start of monitoring in 1971, as well as descendants of recent wave of admixture that began in 1982. As a result, all baboons from Amboseli are admixed [mean = 30 to 37% genome-wide anubis ancestry ± 10% SD; (Vilgalys et al., 2022). To assess whether hybrid status predicts body size, we used a measure of hybrid score that quantifies genome-wide estimates of anubis ancestry using low coverage local ancestry estimation (Vilgalys et al., 2022; Wall et al., 2016). Our hybrid scores were obtained following methods in Vilgalys, Fogel et al (2022). The resulting hybrid score has a possible range from 0 (pure yellow) to 1 (pure anubis). We only had hybrid scores for about 45% of our study subjects (54 females for shoulder-rump, 56 females for leg and forearm), resulting in a smaller dataset than our main analyses. To test for a possible effect of hybrid score on body size, we used these smaller datasets to model the effect of the proportion of drought days in the first four years of life on leg length, forearm length, and shoulder-rump length.

To test whether more admixed individuals were larger or smaller than less admixed individuals, we ran two versions of each model: one that included only a linear term of hybrid score, and a second that also included a quadratic term. While hybrid scores are not always well-predicted by relatedness in this population (because females mate multiply and may produce a series of offspring with disparate hybrid scores), the ASReml-R software indicated singularities in our model, perhaps because hybrid scores and relatedness may be somewhat confounded in our relatively small dataset (Butler et al., 2009). Thus, we were forced to remove the random effect of relatedness from these models to obtain model estimates for the effects of hybrid score. In all models, including hybrid score (without pedigree) resulted in a poorer model fit than models with neither hybrid score nor pedigree (models with linear term only: $\Delta AIC_{shoulder-rump}$ = -2.9, $\Delta AIC_{leg}$ = -5.0, $\Delta AIC_{forearm}$ = -4.8; models with linear and quadratic terms: $\Delta AIC_{shoulder-rump}$ = -0.75, $\Delta AIC_{leg}$ = -2.6, $\Delta AIC_{forearm}$ = -3.7). Due to its poor performance in the models and our inability to include hybrid score along with pedigree, we did not include hybrid score in our main analysis. Specifically, the small size of the data set, the absence of relatedness information in the models, and the fact that including hybrid score reduced model fit, all suggest that our results (Tables S5 and S6) are difficult to interpret at best. A larger dataset will be needed to understand whether admixture has any effect on body size in this population.

**Table S5.** Fixed effects from models that included hybrid score as both linear and quadratic predictors of body size; these models use a much smaller dataset than the models run in the main analysis and did not include a random effect of relatedness; results suggest no effect of hybrid score on body size, but are at best difficult to interpret given the absence of a pedigree in the model.

| Body Part | Predictor | Coefficient | SE | $e^{Coefficient}$ [a] | $e^{SE}$ [a] | P-Value |
|---|---|---|---|---|---|---|
| Shoulder-Rump | Intercept | 3.300 | 0.300 | 27.101 | 1.350 | < 2.22E-16 |
| | log(Age) | 0.124 | 0.278 | 1.132 | 1.321 | 0.656 |
| | $log(Age)^2$ | -0.017 | 0.055 | 0.984 | 1.057 | 0.763 |
| | Maternal Proportional Rank [b] | 0.001 | 0.018 | 1.001 | 1.019 | 0.937 |
| | Drought Days in 4 Years | 0.120 | 0.162 | 1.128 | 1.176 | 0.459 |
| | Hybrid Score | 1.129 | 0.582 | 3.093 | 1.790 | 0.053 |
| | $Hybrid Score^2$ | -1.208 | 0.683 | 0.299 | 1.980 | 0.077 |
| Leg | Intercept | 3.324 | 0.294 | 27.759 | 1.342 | < 2.22E-16 |
| | log(Age) | 0.248 | 0.274 | 1.282 | 1.315 | 0.365 |
| | $log(Age^2)$ | -0.049 | 0.055 | 0.953 | 1.056 | 0.375 |
| | Leg Bent --> Straight | 0.019 | 0.002 | 1.019 | 1.002 | < 2.22E-16 |
| | Maternal Proportional Rank | 0.022 | 0.020 | 1.022 | 1.020 | 0.270 |

| | | | | | | |
|---|---|---|---|---|---|---|
| | Drought Days in 4 Years | -0.134 | 0.165 | 0.875 | 1.179 | 0.416 |
| | Hybrid Score | 1.061 | 0.594 | 2.888 | 1.812 | 0.074 |
| | Hybrid Score$^2$ | -1.311 | 0.698 | 0.269 | 2.010 | 0.060 |
| Forearm | Intercept | 3.157 | 0.370 | 23.489 | 1.447 | < 2.22E-16 |
| | log(Age) | -0.108 | 0.350 | 0.898 | 1.419 | 0.758 |
| | log(Age)$^2$ | 0.026 | 0.070 | 1.026 | 1.072 | 0.712 |
| | Maternal Proportional Rank | 0.030 | 0.024 | 1.030 | 1.025 | 0.222 |
| | Drought Days in 4 Years | -0.102 | 0.204 | 0.903 | 1.226 | 0.617 |
| | Hybrid Score | 0.855 | 0.711 | 2.352 | 2.036 | 0.229 |
| | Hybrid Score$^2$ | -1.032 | 0.834 | 0.356 | 2.303 | 0.216 |

[a] Each one-unit increase in the predictors is associated with a percent change equivalent to the value in these columns. E.g., 1 extra month of drought predicts a shoulder-rump length 99.9% as long, which is equivalent to a decrease in shoulder-rump length of 0.1%.
[b] Maternal proportional rank at study subject's birth

**Table S6.** Random effects of models that included hybrid score as both linear and quadratic predictors of body size; these models use a much smaller dataset than the models run in the main analysis and did not include a random effect of relatedness. Total variance is the sum of all random effect coefficients for a particular model, and proportion of variance for each random effect's coefficient divided by the total variance.

| Drought Days 4 Years | | | | | |
|---|---|---|---|---|---|
| **Body Part** | **Random Effect** | **Coefficient** | **SE** | **Total Variance** | **Proportion of Variance** |
| Shoulder-rump | Maternal Effect | 6.99E-04 | 3.36E-04 | 2.12E-03 | 0.33 |
| | ID (Between-Individual Variation) | 5.42E-04 | 2.42E-04 | | 0.26 |
| | Residual Variance | 8.80E-04 | 5.41E-05 | | 0.41 |
| Leg | Maternal Effect | 1.04E-03 | 4.12E-04 | 1.94E-03 | 0.53 |
| | ID (Between-Individual Variation) | 5.60E-04 | 2.36E-04 | | 0.29 |
| | Residual Variance | 3.45E-04 | 1.85E-05 | | 0.18 |
| Forearm | Maternal Effect | 1.51E-03 | 6.07E-04 | 3.80E-03 | 0.40 |
| | ID (Between-Individual Variation) | 7.19E-04 | 3.42E-04 | | 0.19 |
| | Residual Variance | 1.57E-03 | 8.12E-05 | | 0.41 |

### Final dataset and model results

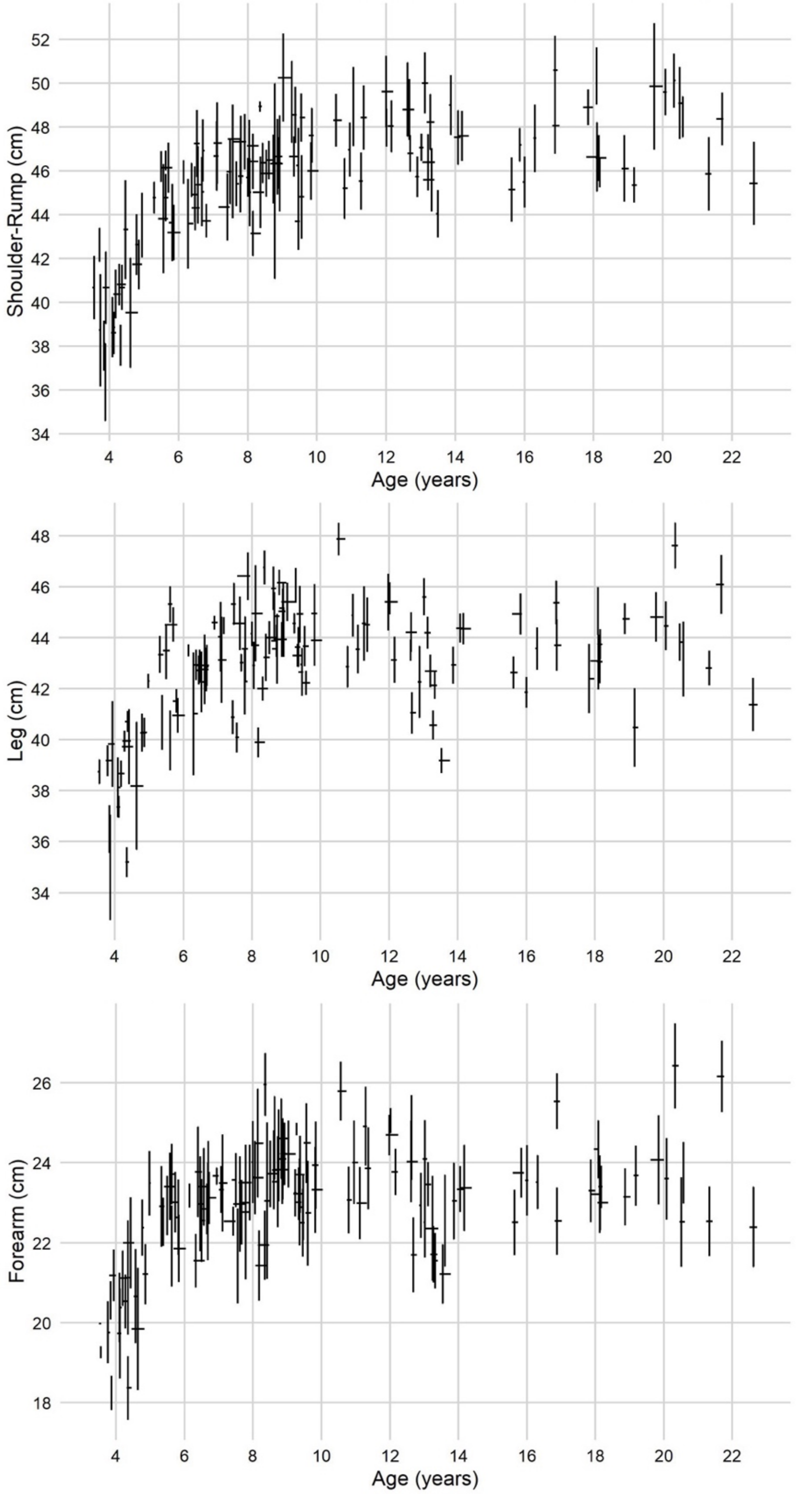

**Figure S4.** Representation of the error in the three body size datasets. Each cross represents one study subject. Vertical error bars are +/- SD. Horizontal bars span the ages over which that subject's data were collected.

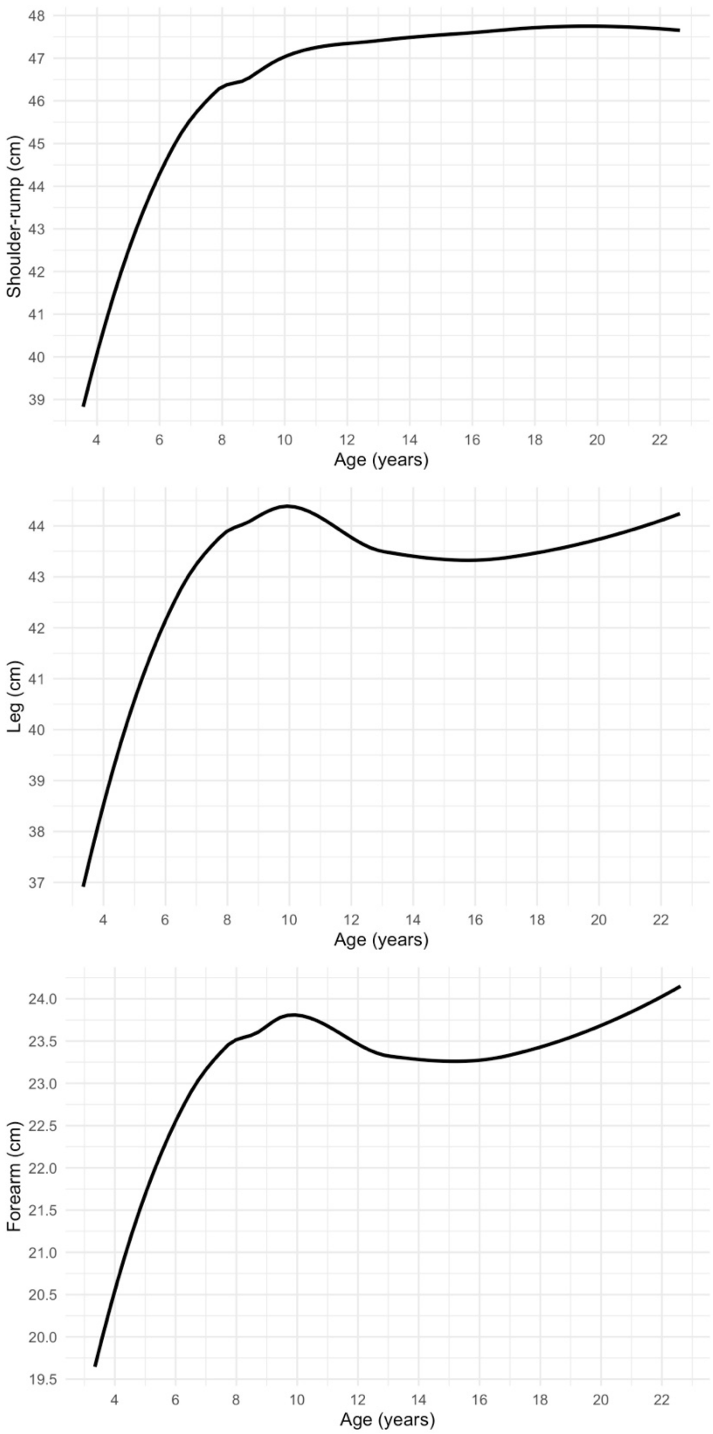

**Figure S5.** Lowess plots of the three body size datasets. Lowess curves were drawn based on the mean body size for each baboon. Curves were drawn using the lowess function in ggplot2's geom_smooth option using the automatic weighting value.

*Cumulative rainfall results*

Cumulative rainfall in the first 4 years of life, but not in the first year of life alone, predicted one measure of body size: leg length (Table S7). Cumulative rainfall did not predict shoulder-rump length or forearm length (Table S7). For leg length, each additional millimeter of rain in the first 4 years of life predicted a 0.005% increase in leg length in juvenescence and adulthood ($p = 0.032$, Table S5). A female who experienced cumulative rainfall that was 1 SD (214 mm) below the mean of 1368 mm rain in the first 4 years of life is predicted to have legs 1.1% shorter. For our study subjects, the amount of rainfall experienced in the first 4 years of life ranged from 1072 to 1818 mm, indicating a 3.7% difference in leg length between the most- and least-affected female.

**Table S7.** Results from the 18 models run to assess effects of early-life environment on body size (6 predictors of early-life environment assessed for 3 body size measures).

| Early-Life Adversity Score | | | | | | | |
|---|---|---|---|---|---|---|---|
| **Body Part** | **Predictor** | **Coefficient** | **SE** | **$e^{Coefficient}$ a** | **$e^{SE}$ a** | **P-Value** | **Interpretation** |
| Shoulder-Rump | Intercept | 3.072 | 0.059 | 21.589 | 1.061 | < 2.22E-16 | At birth, baboons are predicted to have a shoulder-rump length of 21.6 cm |
| | log(Age) | 0.609 | 0.056 | 1.839 | 1.058 | < 2.22E-16 | Shoulder-rump length is strongly predicted by age |
| | $log(Age)^2$ | -0.116 | 0.013 | 0.890 | 1.013 | < 2.22E-16 | |
| | Early-Life Adversity Score | -0.001 | 0.003 | 0.999 | 1.003 | 0.734 | No effect of early-life adversity score on shoulder-rump length |
| Leg | Intercept | 2.992 | 0.063 | 19.919 | 1.065 | < 2.22E-16 | At birth, baboons are predicted to have a leg length of 19.7 cm |
| | log(Age) | 0.629 | 0.061 | 1.875 | 1.063 | < 2.22E-16 | Leg length is strongly predicted by age |
| | $log(Age^2)$ | -0.127 | 0.014 | 0.881 | 1.014 | < 2.22E-16 | |
| | Leg Bent --> Straight | 0.017 | 0.001 | 1.017 | 1.001 | < 2.22E-16 | Images in which the leg was straight have leg lengths 1.7% larger than images in which the leg was slightly bent |
| | Early-Life Adversity Score | -0.003 | 0.004 | 0.997 | 1.004 | 0.398 | No effect of early-life adversity score on leg length |
| Forearm | Intercept | 2.442 | 0.074 | 11.490 | 1.077 | < 2.22E-16 | At birth, baboons are predicted to have a forearm length of 11.3 cm |
| | log(Age) | 0.580 | 0.071 | 1.785 | 1.073 | 3.33E-16 | Forearm length is strongly predicted by age |
| | $log(Age)^2$ | -0.115 | 0.016 | 0.891 | 1.016 | 8.91E-13 | |
| | Early-Life Adversity Score | 0.001 | 0.004 | 1.001 | 1.004 | 0.896 | No effect of early-life adversity score on forearm length |
| **Maternal Death** | | | | | | | |
| **Body Part** | **Predictor** | **Coefficient** | **SE** | **$e^{Coefficient}$ 1** | **$e^{SE}$ 1** | **P-Value** | **Interpretation** |
| Shoulder-Rump | Intercept | 3.075 | 0.058 | 21.640 | 1.060 | < 2.22E-16 | At birth, baboons are predicted to have a shoulder-rump length of 21.6 cm |
| | log(Age) | 0.608 | 0.053 | 1.836 | 1.055 | < 2.22E-16 | Shoulder-rump length is strongly predicted by age |
| | $log(Age)^2$ | -0.116 | 0.012 | 0.890 | 1.012 | < 2.22E-16 | |

| | | | | | | | |
|---|---|---|---|---|---|---|---|
| | Maternal Proportional Rank [b] | -0.001 | 0.012 | 0.999 | 1.012 | 0.917 | No effect of maternal rank on shoulder-rump length |
| | Maternal Death | 0.007 | 0.007 | 1.007 | 1.008 | 0.333 | No effect of maternal death on shoulder-rump length |
| Leg | Intercept | 2.999 | 0.061 | 20.070 | 1.063 | < 2.22E-16 | At birth, baboons are predicted to have a leg length of 20.1 cm |
| | log(Age) | 0.608 | 0.057 | 1.837 | 1.059 | < 2.22E-16 | Leg length is strongly predicted by age |
| | log(Age)$^2$ | -0.122 | 0.013 | 0.885 | 1.013 | < 2.22E-16 | |
| | Leg Bent --> Straight | 0.017 | 0.001 | 1.017 | 1.001 | < 2.22E-16 | Images in which the leg was straight have leg lengths 1.7% larger than images in which the leg was slightly bent |
| | Maternal Proportional Rank | 0.021 | 0.014 | 1.021 | 1.014 | 0.140 | Very weak pattern: The daughter of the highest-ranking mother is predicted to have legs that are 2.1% longer than the daughter of the lowest-ranking mother |
| | Maternal Death | -0.002 | 0.008 | 0.998 | 1.008 | 0.837 | No effect of maternal death on leg length |
| Forearm | Intercept | 2.418 | 0.072 | 11.228 | 1.074 | < 2.22E-16 | At birth, baboons are predicted to have a forearm length of 11.2 cm |
| | log(Age) | 0.588 | 0.066 | 1.801 | 1.069 | < 2.22E-16 | Forearm length is strongly predicted by age |
| | log(Age)$^2$ | -0.117 | 0.015 | 0.890 | 1.015 | 8.66E-15 | |
| | Maternal Proportional Rank | 0.027 | 0.016 | 1.027 | 1.016 | 0.088 | Trending: The daughter of the highest-ranking mother is predicted to have forearms that are 2.7% longer than the daughter of the lowest-ranking mother |
| | Maternal Death | 0.007 | 0.009 | 1.007 | 1.009 | 0.476 | No effect of maternal death on forearm length |

**Rainfall 1 Year**

| Body Part | Predictor | Coefficient | SE | $e^{\text{Coefficient}}$ [1] | $e^{\text{SE}}$ [1] | P-Value | Interpretation |
|---|---|---|---|---|---|---|---|
| Shoulder-Rump | Intercept | 3.092 | 0.066 | 22.017 | 1.069 | < 2.22E-16 | At birth, baboons are predicted to have a shoulder-rump length of 22.0 cm |
| | log(Age) | 0.598 | 0.057 | 1.819 | 1.058 | < 2.22E-16 | Shoulder-rump length is strongly predicted by age |
| | log(Age)$^2$ | -0.114 | 0.013 | 0.892 | 1.013 | < 2.22E-16 | |
| | Maternal Proportional Rank | -0.001 | 0.012 | 0.999 | 1.012 | 0.953 | No effect of maternal rank on shoulder-rump length |

| | | | | | | | |
|---|---|---|---|---|---|---|---|
| | Rainfall in 1 Year | -1.72E-05 | 3.00E-05 | 0.99998 | 1.00003 | 0.567 | No effect of rainfall on shoulder-rump length |
| Leg | Intercept | 3.002 | 0.066 | 20.124 | 1.068 | < 2.22E-16 | At birth, baboons are predicted to have a leg length of 20.1 cm |
| | log(Age) | 0.606 | 0.058 | 1.832 | 1.060 | < 2.22E-16 | Leg length is strongly predicted by age |
| | $\log(\text{Age})^2$ | -0.122 | 0.013 | 0.885 | 1.013 | < 2.22E-16 | |
| | Leg Bent --> Straight | 0.017 | 0.001 | 1.017 | 1.001 | < 2.22E-16 | Images in which the leg was straight have leg lengths 1.7% larger than images in which the leg was slightly bent |
| | Maternal Proportional Rank | 0.020 | 0.014 | 1.021 | 1.014 | 0.142 | Very weak pattern: The daughter of the highest-ranking mother is predicted to have legs that are 2.1% longer than the daughter of the lowest-ranking mother |
| | Rainfall in 1 Year | -2.86E-06 | 3.15E-05 | 0.999997 | 1.00003 | 0.927 | No effect of rainfall on leg length |
| Forearm | Intercept | 2.449 | 0.078 | 11.571 | 1.081 | < 2.22E-16 | At birth, baboons are predicted to have a forearm length of 11.6 cm |
| | log(Age) | 0.574 | 0.068 | 1.775 | 1.071 | < 2.22E-16 | Forearm length is strongly predicted by age |
| | $\log(\text{Age})^2$ | -0.114 | 0.015 | 0.892 | 1.016 | 1.64E-13 | |
| | Maternal Proportional Rank | 0.027 | 0.016 | 1.027 | 1.016 | 0.085 | Trending: The daughter of the highest-ranking mother is predicted to have forearms that are 2.7% longer than the daughter of the lowest-ranking mother |
| | Rainfall in 1 Year | -3.44E-05 | 3.58E-05 | 0.99997 | 1.00004 | 0.336 | No effect of rainfall on forearm length |
| **Rainfall 4 Years** | | | | | | | |
| **Body Part** | **Predictor** | **Coefficient** | **SE** | **$e^{\text{Coefficient}}$ [1]** | **$e^{\text{SE}}$ [1]** | **P-Value** | **Interpretation** |
| Shoulder-Rump | Intercept | 3.106 | 0.079 | 22.342 | 1.082 | < 2.22E-16 | At birth, baboons are predicted to have a shoulder-rump length of 22.3 cm |
| | log(Age) | 0.598 | 0.056 | 1.818 | 1.058 | < 2.22E-16 | Shoulder-rump length is strongly predicted by age |
| | $\log(\text{Age})^2$ | -0.114 | 0.012 | 0.892 | 1.012 | < 2.22E-16 | |
| | Maternal Proportional Rank | -4.69E-05 | 0.012 | 1.000 | 1.012 | 0.997 | No effect of maternal rank on shoulder-rump length |
| | Rainfall in 4 Years | -1.33E-05 | 2.16E-05 | 0.999987 | 1.00002 | 0.537 | No effect of rainfall on shoulder-rump length |
| Leg | Intercept | 2.886 | 0.081 | 17.920 | 1.085 | < 2.22E-16 | At birth, baboons are predicted to have a leg length of 17.9 cm |

| | | | | | | | |
|---|---|---|---|---|---|---|---|
| | log(Age) | 0.632 | 0.059 | 1.882 | 1.061 | < 2.22E-16 | Leg length is strongly predicted by age |
| | log(Age)$^2$ | -0.125 | 0.013 | 0.882 | 1.013 | < 2.22E-16 | |
| | Leg Bent --> Straight | 0.017 | 0.001 | 1.017 | 1.001 | < 2.22E-16 | Images in which the leg was straight have leg lengths 1.7% larger than images in which the leg was slightly bent |
| | Maternal Proportional Rank | 0.018 | 0.014 | 1.018 | 1.014 | 0.182 | Very weak pattern: The daughter of the highest-ranking mother is predicted to have legs that are 1.8% longer than the daughter of the lowest-ranking mother |
| | Rainfall in 4 Years | 4.95E-05 | 2.32E-05 | 1.00005 | 1.00002 | 0.032 | Every additional millimeter of rain in the first 4 years of life is predicted to increase later-life leg length by 0.005% |
| Forearm | Intercept | 2.359 | 0.096 | 10.584 | 1.100 | < 2.22E-16 | At birth, baboons are predicted to have a forearm length of 10.6 cm |
| | log(Age) | 0.609 | 0.069 | 1.839 | 1.072 | < 2.22E-16 | Forearm length is strongly predicted by age |
| | log(Age)$^2$ | -0.121 | 0.015 | 0.886 | 1.016 | 5.11E-15 | |
| | Maternal Proportional Rank | 0.026 | 0.016 | 1.027 | 1.016 | 0.095 | Trending: The daughter of the highest-ranking mother is predicted to have forearms that are 2.7% longer than the daughter of the lowest-ranking mother |
| | Rainfall in 4 Years | 2.46E-05 | 2.66E-05 | 1.00002 | 1.00003 | 0.355 | No effect of rainfall on forearm length |
| | | | | **Drought Days 1 Year** | | | |
| **Body Part** | **Predictor** | **Coefficient** | **SE** | **e$^{\text{Coefficient 1}}$** | **e$^{\text{SE 1}}$** | **P-Value** | **Interpretation** |
| Shoulder-Rump | Intercept | 3.073 | 0.058 | 21.604 | 1.060 | < 2.22E-16 | At birth, baboons are predicted to have a shoulder-rump length of 21.6 cm |
| | log(Age) | 0.611 | 0.061 | 1.843 | 1.063 | < 2.22E-16 | Shoulder-rump length is strongly predicted by age |
| | log(Age)$^2$ | -0.117 | 0.014 | 0.890 | 1.014 | < 2.22E-16 | |
| | Maternal Proportional Rank | -0.001 | 0.012 | 0.999 | 1.012 | 0.966 | No effect of maternal rank on shoulder-rump length |
| | Drought Days in 1 Year | -0.002 | 0.039 | 0.998 | 1.040 | 0.952 | No effect of drought on shoulder-rump length |
| Leg | Intercept | 3.005 | 0.061 | 20.183 | 1.063 | < 2.22E-16 | At birth, baboons are predicted to have a leg length of 20.2 cm |
| | log(Age) | 0.627 | 0.062 | 1.872 | 1.064 | < 2.22E-16 | Leg length is strongly predicted by age |
| | log(Age)$^2$ | -0.126 | 0.014 | 0.882 | 1.014 | < 2.22E-16 | |

| | | | | | | | |
|---|---|---|---|---|---|---|---|
| | Leg Bent --> Straight | 0.017 | 0.001 | 1.017 | 1.001 | < 2.22E-16 | Images in which the leg was straight have leg lengths 1.7% larger than images in which the leg was slightly bent |
| | Maternal Proportional Rank | 0.021 | 0.014 | 1.021 | 1.014 | 0.127 | No effect of maternal rank on leg length |
| | Drought Days in 1 Year | -0.041 | 0.042 | 0.960 | 1.042 | 0.327 | No effect of drought on leg length |
| Forearm | Intercept | 2.419 | 0.072 | 11.234 | 1.074 | < 2.22E-16 | At birth, baboons are predicted to have a forearm length of 11.2 cm |
| | log(Age) | 0.602 | 0.072 | 1.825 | 1.075 | 1.11E-16 | Forearm length is strongly predicted by age |
| | log(Age)$^2$ | -0.120 | 0.016 | 0.887 | 1.016 | 1.59E-13 | |
| | Maternal Proportional Rank | 0.028 | 0.016 | 1.028 | 1.016 | 0.080 | Trending: The daughter of the highest-ranking mother is predicted to have forearms that are 2.8% longer than the daughter of the lowest-ranking mother |
| | Drought Days in 1 Year | -0.017 | 0.047 | 0.983 | 1.048 | 0.712 | No effect of drought on forearm length |
| **Drought Days 4 Years** | | | | | | | |
| **Body Part** | **Predictor** | **Coefficient** | **SE** | **e$^{Coefficient}$ [1]** | **e$^{SE}$ [1]** | **P-Value** | **Interpretation** |
| Shoulder-Rump | Intercept | 3.084 | 0.078 | 21.837 | 1.081 | < 2.22E-16 | At birth, baboons are predicted to have a shoulder-rump length of 21.5 cm |
| | log(Age) | 0.612 | 0.054 | 1.844 | 1.056 | < 2.22E-16 | Shoulder-rump length is strongly predicted by age |
| | log(Age)$^2$ | -0.117 | 0.012 | 0.890 | 1.012 | < 2.22E-16 | |
| | Maternal Proportional Rank | -0.001 | 0.012 | 0.999 | 1.012 | 0.960 | No effect of maternal rank on shoulder-rump length |
| | Drought Days in 1 Year | -0.017 | 0.084 | 0.983 | 1.087 | 0.840 | No effect of drought on shoulder-rump length |
| Leg | Intercept | 3.208 | 0.085 | 24.741 | 1.088 | < 2.22E-16 | At birth, baboons are predicted to have a leg length of 24.7 cm |
| | log(Age) | 0.591 | 0.055 | 1.806 | 1.056 | < 2.22E-16 | Leg length is strongly predicted by age |
| | log(Age)$^2$ | -0.117 | 0.013 | 0.889 | 1.013 | < 2.22E-16 | |
| | Leg Bent --> Straight | 0.017 | 0.001 | 1.017 | 1.001 | < 2.22E-16 | Images in which the leg was straight have leg lengths 1.7% larger than images in which the leg was slightly bent |

| | | | | | | | |
|---|---|---|---|---|---|---|---|
| | Maternal Proportional Rank | 0.018 | 0.013 | 1.019 | 1.013 | 0.163 | No effect of maternal rank on leg length |
| | Drought Days in 4 Years | -0.262 | 0.089 | 0.770 | 1.093 | 0.003 | A 1% increase in proportion of drought days predicts a decrease in leg length of 0.23%; each additional day of drought predicts a decrease in leg length of 0.015% |
| Forearm | Intercept | 2.631 | 0.098 | 13.891 | 1.103 | < 2.22E-16 | At birth, baboons are predicted to have a forearm length of 13.9 cm |
| | log(Age) | 0.598 | 0.063 | 1.818 | 1.065 | < 2.22E-16 | Forearm length is strongly predicted by age |
| | $log(Age)^2$ | -0.118 | 0.014 | 0.889 | 1.014 | 1.11E-16 | |
| | Maternal Proportional Rank | 0.026 | 0.015 | 1.027 | 1.015 | 0.083 | Trending: The daughter of the highest-ranking mother is predicted to have forearms that are 2.7% longer than the daughter of the lowest-ranking mother |
| | Drought Days in 1 Year | -0.286 | 0.102 | 0.751 | 1.107 | 0.005 | A 1% increase in proportion of drought days predicts a decrease in leg length of 0.25%; each additional day of drought predicts a decrease in leg length of 0.018% |

[a] Every one-unit increase in the predictors is associated with a percent change equivalent to the value in these columns. E.g., 1 extra month of drought predicts a shoulder-rump length 99.9% as long, which is equivalent to a decrease in shoulder-rump length of 0.1%.
[b] Maternal proportional rank at study subject's birth

**Table S8.** Random effects from all 18 models run to assess effects of early-life environment on body size (6 predictors of early-life environment assessed for 3 body size measures); total variance is the sum of all random effect coefficients for a particular model, and proportion of variance for each random effect's coefficient divided by the total variance.

| Body Part | Random Effect | Coefficient | SE [a] | Total Variance | Proportion of Variance |
|---|---|---|---|---|---|
| **Early-Life Adversity Score** | | | | | |
| Shoulder-rump | Additive Genetic Effect | 1.27E-03 | 1.93E-04 | 2.20E-03 | 0.58 |
| | Maternal Effect | 5.33E-10 | NA | | 2.42E-07 |
| | ID (Between-Individual Variation) | 2.11E-10 | NA | | 9.58E-08 |
| | Residual Variance | 9.29E-04 | 4.23E-05 | | 0.42 |
| Leg | Additive Genetic Effect | 1.33E-03 | 2.66E-04 | 2.10E-03 | 0.63 |
| | Maternal Effect | 3.44E-04 | 2.22E-04 | | 0.16 |
| | ID (Between-Individual Variation) | 2.91E-07 | 2.65E-06 | | 1.38E-04 |
| | Residual Variance | 4.30E-04 | 1.69E-05 | | 0.20 |
| Forearm | Additive Genetic Effect | 1.48E-03 | 3.25E-04 | 3.69E-03 | 0.40 |
| | Maternal Effect | 5.17E-04 | 2.87E-04 | | 0.14 |
| | ID (Between-Individual Variation) | 1.32E-10 | NA | | 3.58E-08 |
| | Residual Variance | 1.69E-03 | 6.42E-05 | | 0.46 |
| **Maternal Death** | | | | | |
| **Body Part** | **Random Effect** | **Coefficient** | **SE [a]** | **Total Variance** | **Proportion of Variance** |
| Shoulder-rump | Additive Genetic Effect | 1.27E-03 | 1.93E-04 | 2.20E-03 | 0.58 |
| | Maternal Effect | 5.33E-10 | NA | | 2.42E-07 |
| | ID (Between-Individual Variation) | 2.11E-10 | NA | | 9.58E-08 |
| | Residual Variance | 9.29E-04 | 4.23E-05 | | 0.42 |
| Leg | Additive Genetic Effect | 1.33E-03 | 2.66E-04 | 2.10E-03 | 0.63 |
| | Maternal Effect | 3.44E-04 | 2.22E-04 | | 0.16 |
| | ID (Between-Individual Variation) | 2.91E-07 | 2.65E-06 | | 1.38E-04 |
| | Residual Variance | 4.30E-04 | 1.69E-05 | | 0.20 |
| Forearm | Additive Genetic Effect | 1.48E-03 | 3.25E-04 | 3.69E-03 | 0.40 |
| | Maternal Effect | 5.17E-04 | 2.87E-04 | | 0.14 |
| | ID (Between-Individual Variation) | 1.32E-10 | NA | | 3.58E-08 |
| | Residual Variance | 1.69E-03 | 6.42E-05 | | 0.46 |
| **Rainfall 1 Year** | | | | | |
| **Body Part** | **Random Effect** | **Coefficient** | **SE [a]** | **Total Variance** | **Proportion of Variance** |
| Shoulder-rump | Additive Genetic Effect | 1.27E-03 | 1.94E-04 | 2.20E-03 | 0.58 |
| | Maternal Effect | 3.98E-10 | NA | | 1.81E-07 |
| | ID (Between-Individual Variation) | 5.02E-11 | NA | | 2.28E-08 |
| | Residual Variance | 9.29E-04 | 4.24E-05 | | 0.42 |
| Leg | Additive Genetic Effect | 1.33E-03 | 2.64E-04 | 2.10E-03 | 0.63 |
| | Maternal Effect | 3.47E-04 | 2.20E-04 | | 0.17 |

| | | | | | |
|---|---|---|---|---|---|
| | ID (Between-Individual Variation) | 3.63E-07 | 2.69E-06 | | 1.72E-04 |
| | Residual Variance | 4.30E-04 | 1.69E-05 | | 0.20 |
| Forearm | Additive Genetic Effect | 1.50E-03 | 3.29E-04 | 3.67E-03 | 0.41 |
| | Maternal Effect | 4.76E-04 | 2.82E-04 | | 0.13 |
| | ID (Between-Individual Variation) | 1.05E-10 | NA | | 2.86E-08 |
| | Residual Variance | 1.69E-03 | 6.42E-05 | | 0.46 |
| **Rainfall 4 Years** | | | | | |
| **Body Part** | **Random Effect** | **Coefficient** | **SE** [a] | **Total Variance** | **Proportion of Variance** |
| Shoulder-rump | Additive Genetic Effect | 1.27E-03 | 1.93E-04 | 2.20E-03 | 0.58 |
| | Maternal Effect | 5.86E-10 | NA | | 2.66E-07 |
| | ID (Between-Individual Variation) | 1.19E-10 | NA | | 5.41E-08 |
| | Residual Variance | 9.29E-04 | 4.23E-05 | | 0.42 |
| Leg | Additive Genetic Effect | 1.33E-03 | 2.64E-04 | 2.06E-03 | 0.65 |
| | Maternal Effect | 2.93E-04 | 2.09E-04 | | 0.14 |
| | ID (Between-Individual Variation) | 1.73E-06 | 4.54E-06 | | 8.40E-04 |
| | Residual Variance | 4.29E-04 | 1.69E-05 | | 0.21 |
| Forearm | Additive Genetic Effect | 1.53E-03 | 3.33E-04 | 3.68E-03 | 0.41 |
| | Maternal Effect | 4.65E-04 | 2.82E-04 | | 0.13 |
| | ID (Between-Individual Variation) | 8.25E-10 | NA | | 2.24E-07 |
| | Residual Variance | 1.69E-03 | 6.41E-05 | | 0.46 |
| **Drought Days 1 Year** | | | | | |
| **Body Part** | **Random Effect** | **Coefficient** | **SE** [a] | **Total Variance** | **Proportion of Variance** |
| Shoulder-rump | Additive Genetic Effect | 1.28E-03 | 1.95E-04 | 2.21E-03 | 0.58 |
| | Maternal Effect | 5.35E-10 | NA | | 2.42E-07 |
| | ID (Between-Individual Variation) | 1.38E-10 | NA | | 6.26E-08 |
| | Residual Variance | 9.29E-04 | 4.23E-05 | | 0.42 |
| Leg | Additive Genetic Effect | 1.32E-03 | 2.63E-04 | 2.10E-03 | 0.63 |
| | Maternal Effect | 3.50E-04 | 2.18E-04 | | 0.17 |
| | ID (Between-Individual Variation) | 8.00E-07 | 3.26E-06 | | 3.81E-04 |
| | Residual Variance | 4.29E-04 | 1.69E-05 | | 0.20 |
| Forearm | Additive Genetic Effect | 1.51E-03 | 3.30E-04 | 3.69E-03 | 0.41 |
| | Maternal Effect | 4.95E-04 | 2.86E-04 | | 0.13 |
| | ID (Between-Individual Variation) | 9.66E-10 | NA | | 2.62E-07 |
| | Residual Variance | 1.69E-03 | 6.41E-05 | | 0.46 |
| **Drought Days 4 Years** | | | | | |
| **Body Part** | **Random Effect** | **Coefficient** | **SE** [a] | **Total Variance** | **Proportion of Variance** |
| Shoulder-rump | Additive Genetic Effect | 1.28E-03 | 1.94E-04 | 2.21E-03 | 0.58 |
| | Maternal Effect | 5.68E-10 | NA | | 2.57E-07 |

| | | | | | |
|---|---|---|---|---|---|
| | ID (Between-Individual Variation) | 1.34E-10 | NA | | 6.07E-08 |
| | Residual Variance | 9.29E-04 | 4.23E-05 | | 0.42 |
| Leg | Additive Genetic Effect | 1.29E-03 | 2.54E-04 | 1.97E-03 | 0.66 |
| | Maternal Effect | 2.45E-04 | 1.93E-04 | | 0.12 |
| | ID (Between-Individual Variation) | 2.28E-06 | 5.21E-06 | | 1.16E-03 |
| | Residual Variance | 4.30E-04 | 1.69E-05 | | 0.22 |
| Forearm | Additive Genetic Effect | 1.38E-03 | 3.08E-04 | 3.54E-03 | 0.39 |
| | Maternal Effect | 4.76E-04 | 2.72E-04 | | 0.13 |
| | ID (Between-Individual Variation) | 1.71E-10 | NA | | 4.83E-08 |
| | Residual Variance | 1.69E-03 | 6.41E-05 | | 0.48 |

[a] Some components have NA for the standard error because the component coefficients were too close to zero for the model to accurately calculate error.

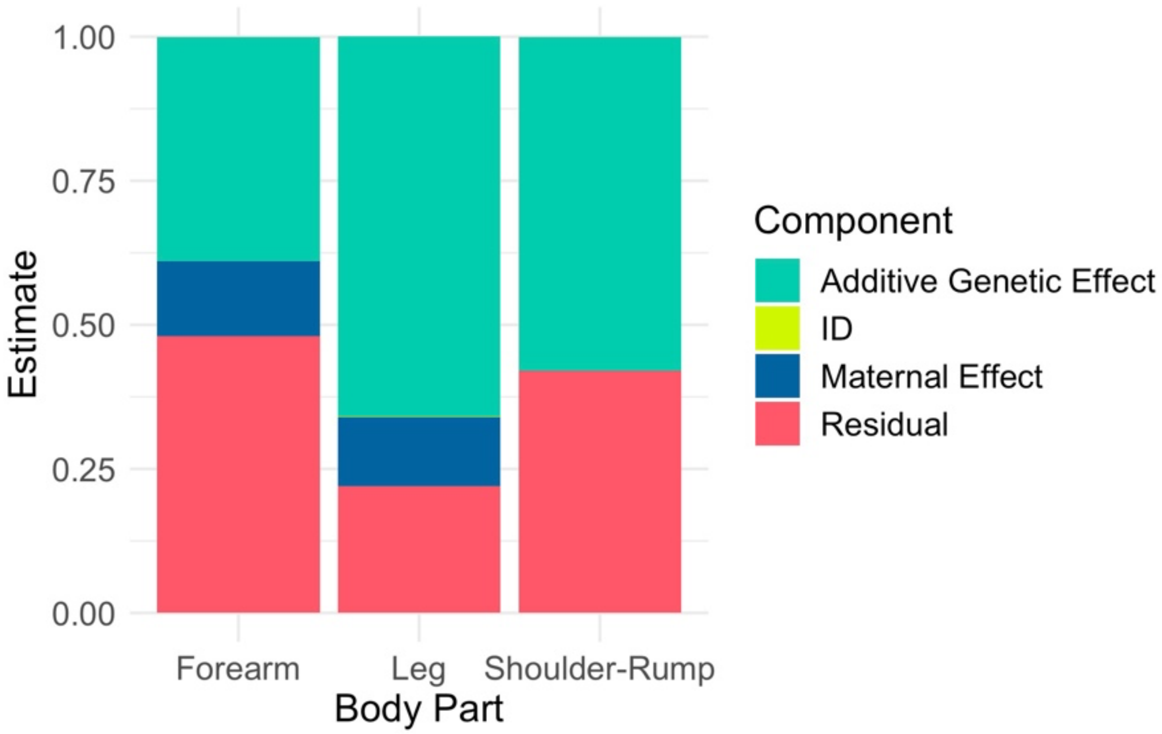

**Figure S6.** Proportion of variance explained by each random effect in three representative models: number of drought days in the first four years of life as a function of (1) forearm length, (2) leg length, and (3) shoulder-rump length.

**Protocol for measuring photogrammetry images**

This document details instructions to take 4 body size measurements: shoulder-rump, femur, lower leg (fibula/tibia), and forearm (radius/ulna). The shoulder-rump measurement also produces a torso height measure. The femur and lower leg measurements will be added together for a leg length measure.

Sophia Li and Emily Levy created the first version of this document in 2017; Levy continues to update it.

Document structure:

- Image set-up & laser distance measurement
- Body size measurements
    - Instructions
    - Figures
    - Examples
        - Shoulder-rump
        - Lower Leg
        - Femur
        - Forearm

*ImageJ set-up & laser distance measurement*

1. Open image in ImageJ by drag an image file into the ImageJ box.
2. A few useful shortcuts:
    a. To measure, press *m*
    b. To zoom in, press *command*+; to zoom out, press *command-*
    c. To scroll sideways (if you're using a mouse instead of a trackpad), press *space bar* while scrolling.
3. Set the measures you want ImageJ to make when you press M.
    a. Analyze → Set measurements → Check Centroid, Bounding Rectangle, and Display Label.
    b. You should only have to do this once – when you re-open ImageJ in the future it will remember your selections.
4. Straighten image
    a. Using line tool, draw a line along the spine. Measure (*m*)
    b. In top bar, go to Image → Transform → Rotate. Enter the angle. Once you click ok, the spine should look like it's horizontal.
    c. To think about what angle to set as the spine, imagine you're creating the top line to measure this baboon's torso. You want to take most of the spine into account, but if the shoulders stick up a lot you don't want to include those.

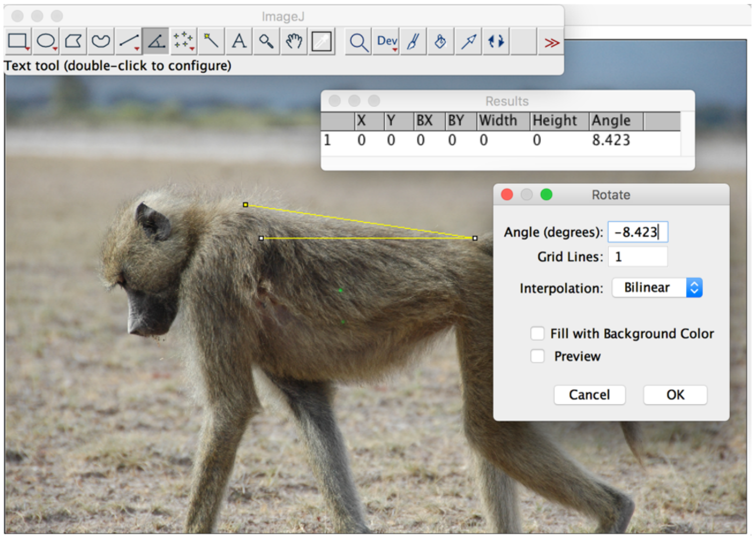

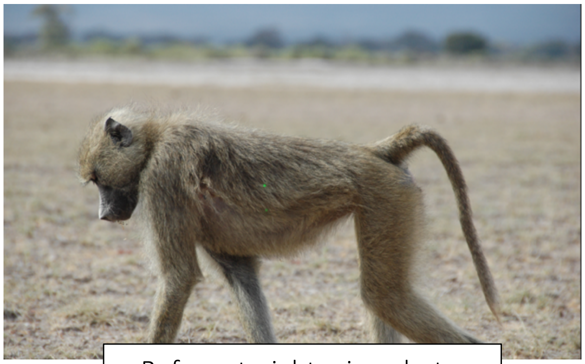

Before straightening photo

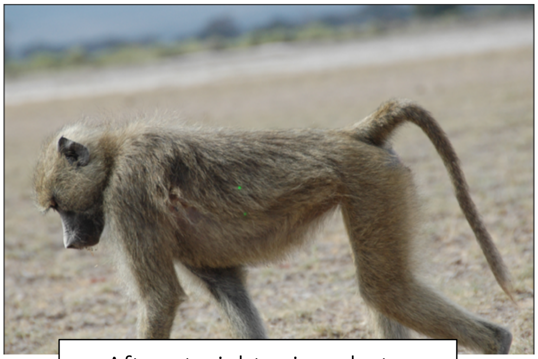

After straightening photo

5. Measure centroid of laser points using oval selection tool

   *Note: We have automated this process, so it's not necessary to do this unless the automated method did not work. For the automated method, see the ImageJ documents here:* [*github.com/ejlevy/Photogrammetry_Coding_InterLaser_Distance*](github.com/ejlevy/Photogrammetry_Coding_InterLaser_Distance)

   a. Select the Oval tool. Draw a circle around one laser.
   b. For most photos, there won't be a really well-defined circle. To help, to go Image → Adjust → Brightness/Contrast. With your laserbeam encircled, press *Auto*. This will adjust the brightness and contrast of the image to accentuate the green of the lasers.
   c. With this coloration, draw a circle or oval around each laser and measure (*m*). You can *Reset* the colors and then press *Auto* again a few times throughout if you want to make sure you're on the mark.
   d. The centroid values (X,Y) are what you will use in analysis (NOT BX, BY). The centroid is just calculating the center of the oval you draw.

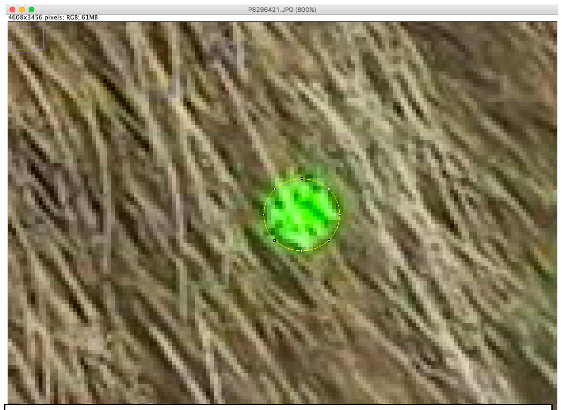

Before adjusting brightness/contrast.

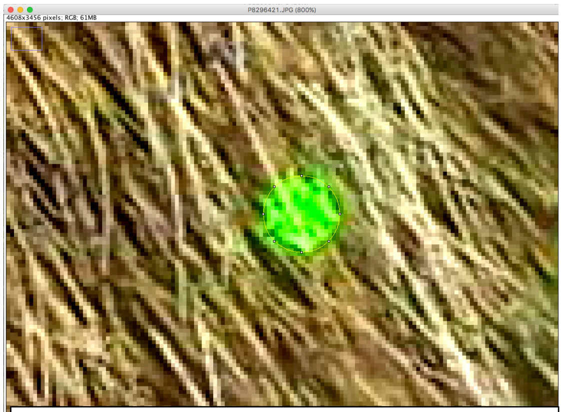

After adjusting brightness/contrast. If lasers are already well-defined, this step won't help much.

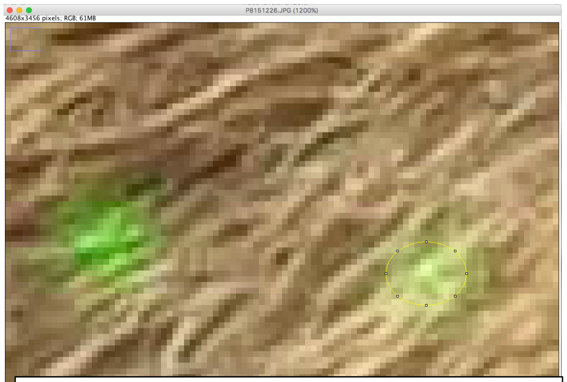

Before adjusting brightness/contrast.

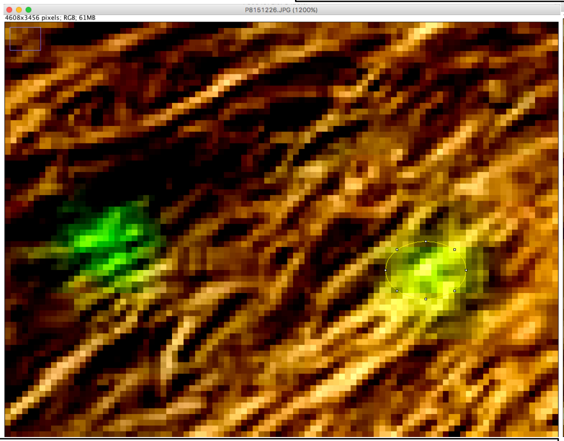

After adjusting brightness/contrast while selection encircles right laser beam.

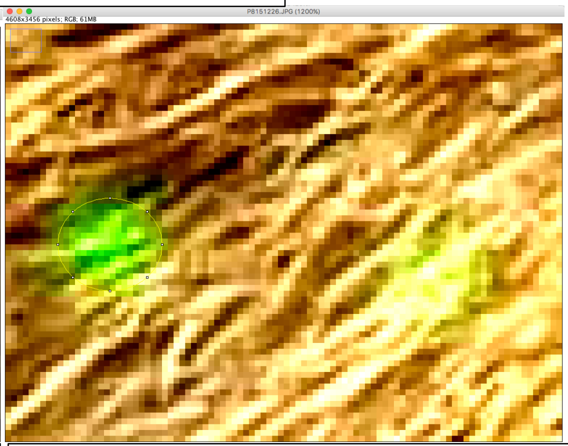

After adjusting brightness/contrast while selection encircles left laser beam.

*Body size measurements*

Instructions

1. Measure shoulder-rump length
    a. Readjust Brightness/Contrast if necessary by pressing *Reset* on Brightness/Contrast panel.
    b. Use rectangle selection tool to draw a rectangle with length from shoulder to rump.
    c. Shoulder is the outer-most point of the shoulder, not including fur.
        i. This measurement can be tricky, especially when the edge of the shoulder blurs into the neck. In that case, you can usually see a difference in the plane of the photo, as the shoulder should be closer to you than the neck (Figures 1, red arrow).
        ii. Looking at a few different photos taken in close succession of the one you're measuring can help.
    d. Rump is the posterior-most point of the ischial callosity (Figure 1, blue arrow).
    e. Press *m*
2. Measure lower leg and femur
    a. Use the line tool to draw a line from the end of the tarsus to the 'knee-pit'.
        i. For the tarsus point, identify the outer-most point on the back of the heel. Think of the back of the heel as a semi-circle, and choose the point at which the curvature from the heel ends (Figure 2, blue arrow).
        ii. For the knee-pit, imagine drawing a line parallel to the shin. For where to stop, imagine two straight lines, one from the ankle to the knee, and a second from the knee to the callosities. Your point should go where those two lines intersect, and should be where the fur still exists. In other words, do not go as posterior as possible looking for a skin boundary (Figure 2, red arrow).
    b. Press *m*
    c. Drag the heel landmark up to the top of the callosity. *Keep the knee-pit point exactly where it was*.
        i. The callosity point should go to the top and posterior-most point of the callosity, not where you see color change (Figure 3).
    d. Press *m*
3. Measure forearm
    a. Use the line tool to draw a line from the elbow to the wrist. Only use the arm closer to the camera.
        i. The elbow is usually occluded by a tuft of fur. The joint itself is usually above and anterior to the longest fur in this tuft (Figures 4 and 5).
        ii. The wrist landmark will be just below the pad (Figure 5).
    b. Press *m*
4. Copy measurements to excel file
    a. Select all measurement and copy into excel.
    b. Label each row with the corresponding measurement
        i. Angle=1, left laser=2, right laser=3, shoulder-rump=4, lower leg=5, femur=6, forearm=7

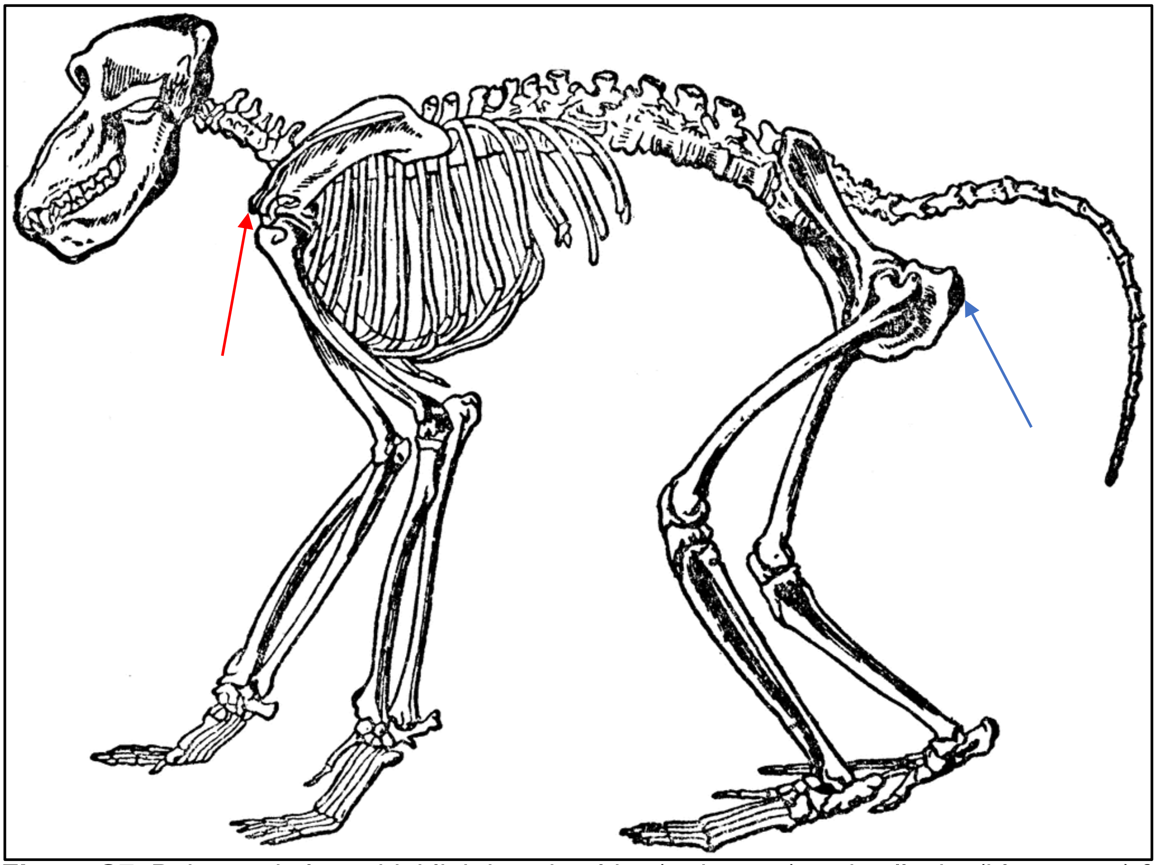

**Figure S7**. Baboon skeleton highlighting shoulder (red arrow) and callosity (blue arrow) for shoulder-rump measure.

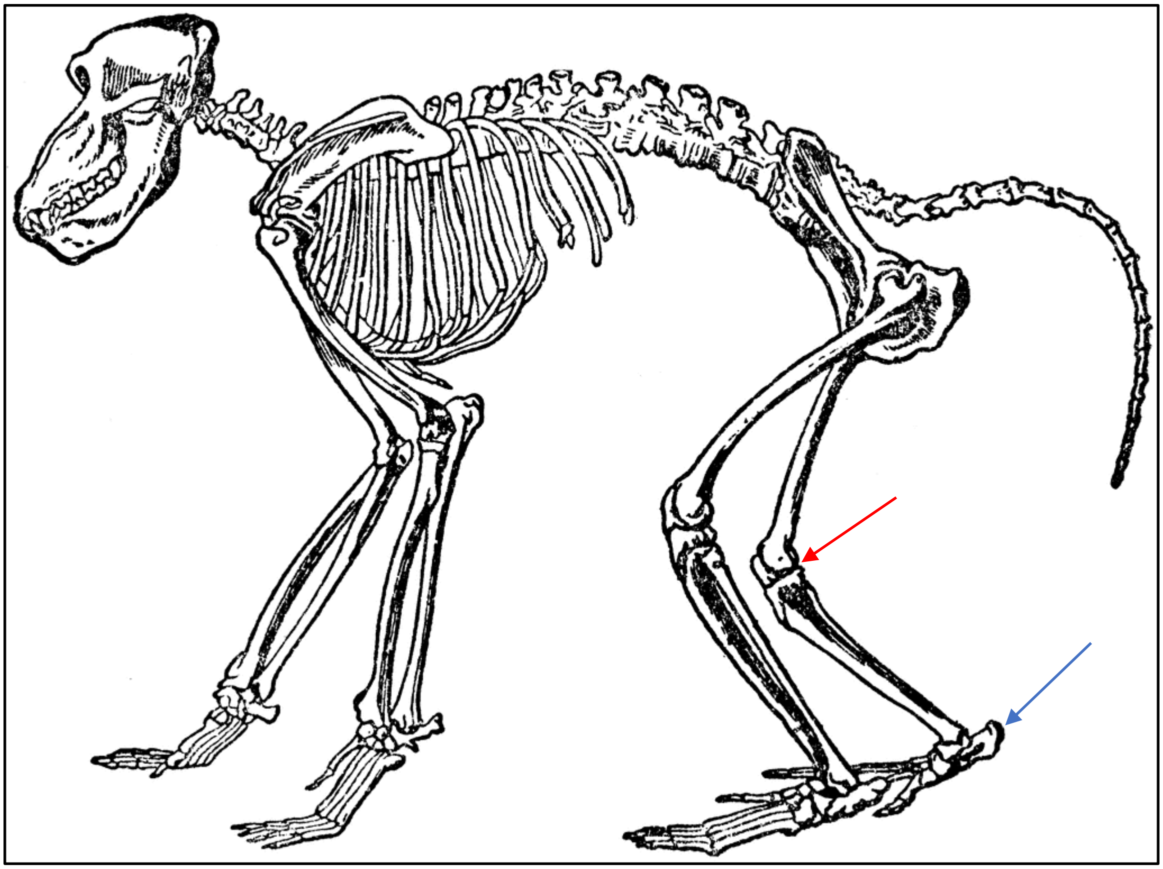

**Figure S8.** Baboon skeleton highlighting knee (red arrow) and heel (blue arrow) for lower leg length measure.

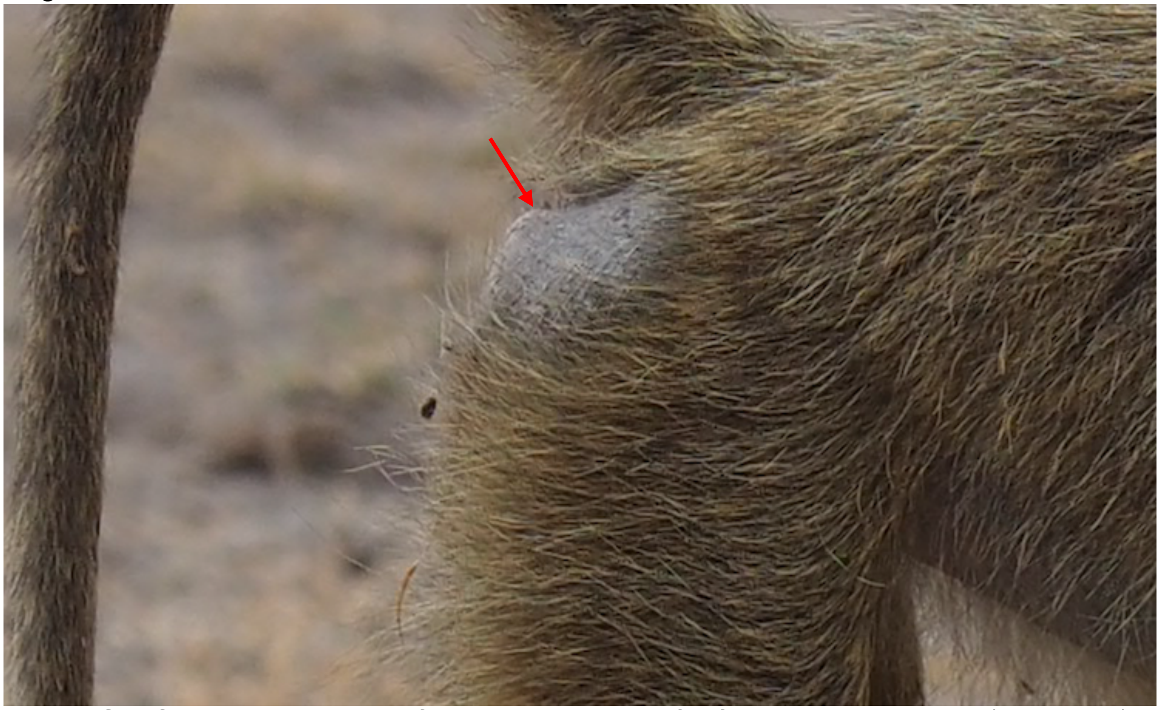

**Figure S9.** Correct placement of callosity landmark for femur measurement (red arrow).

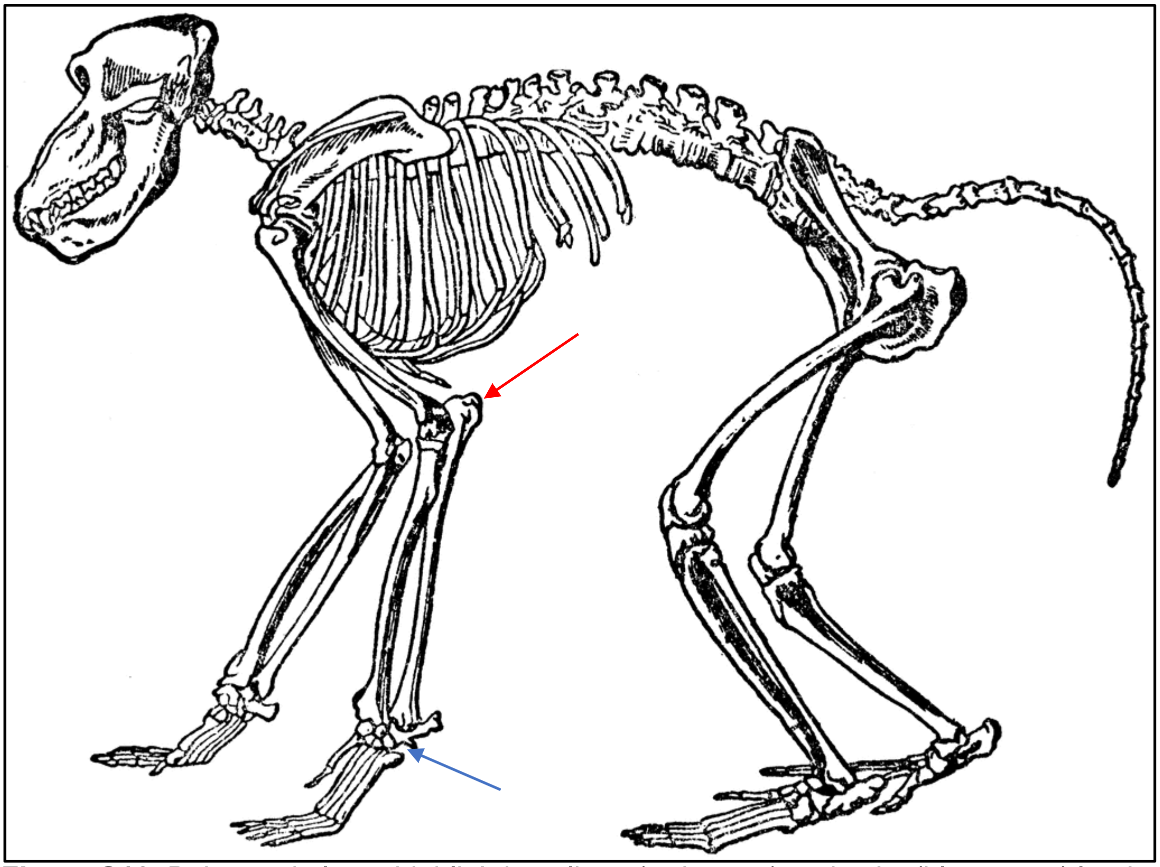

**Figure S10.** Baboon skeleton highlighting elbow (red arrow) and wrist (blue arrow) landmarks for forearm measurement.

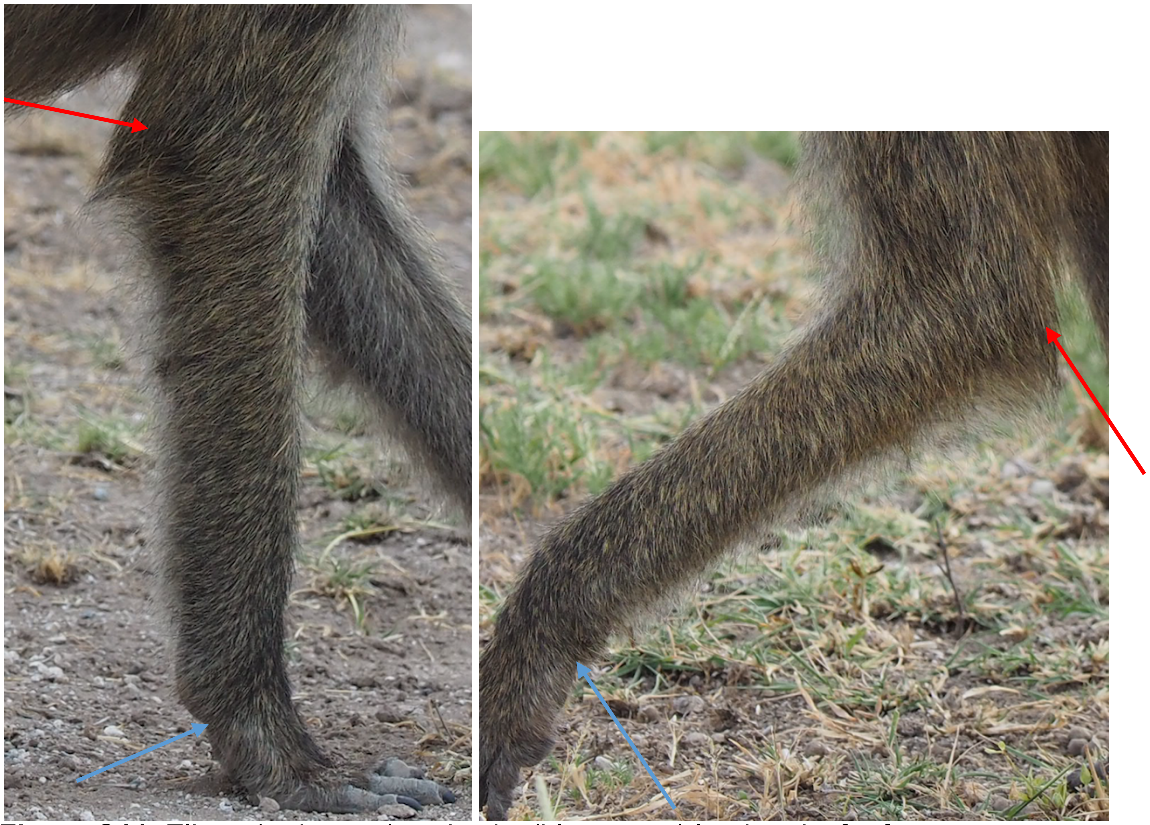

**Figure S11.** Elbow (red arrow) and wrist (blue arrow) landmarks for forearm measurement. Left: Arm straight. Right: Arm bent.

Examples

SHOULDER-RUMP

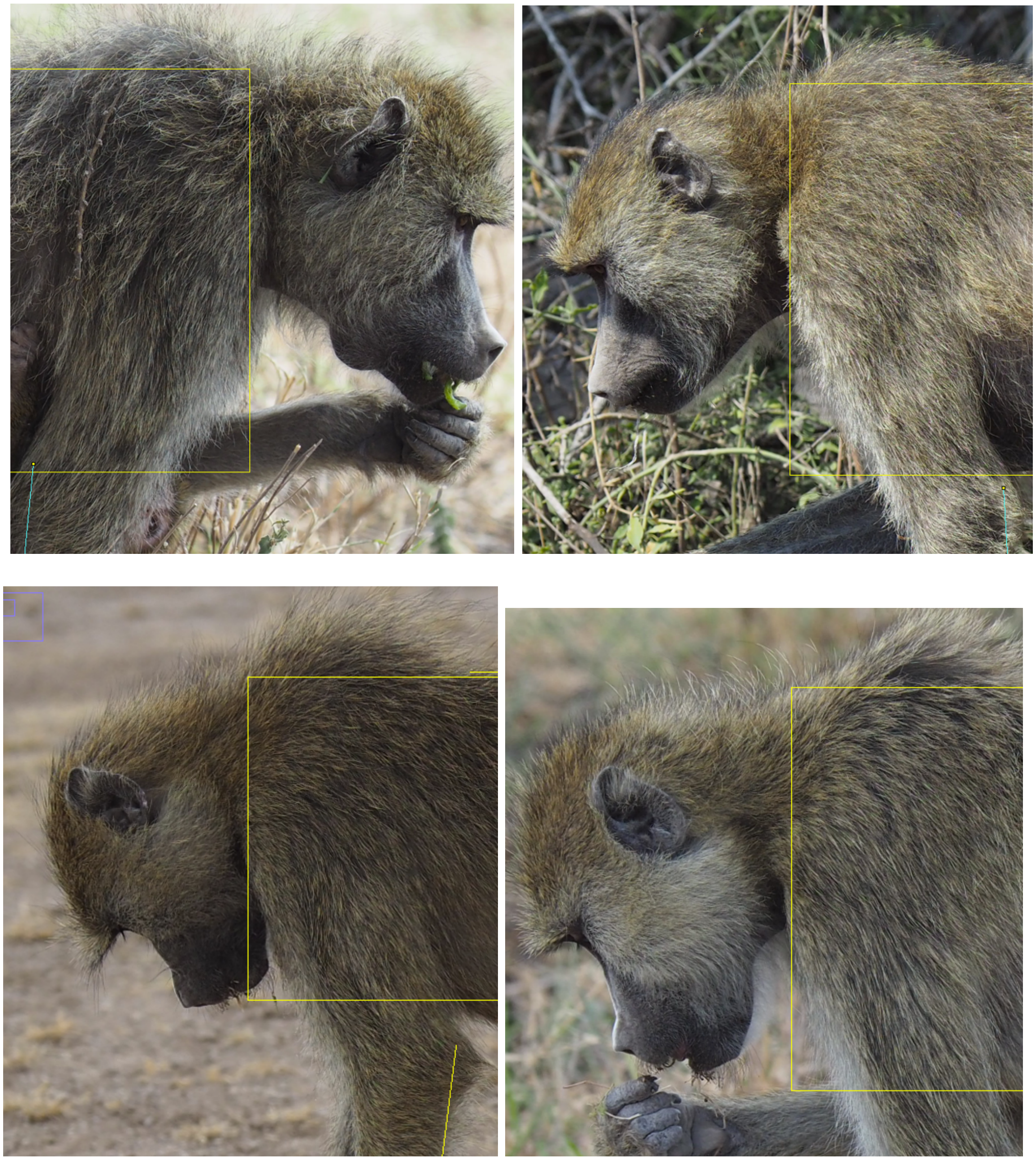

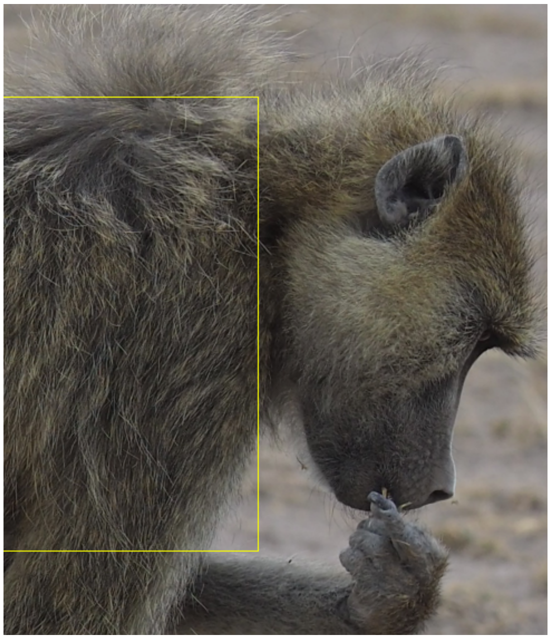
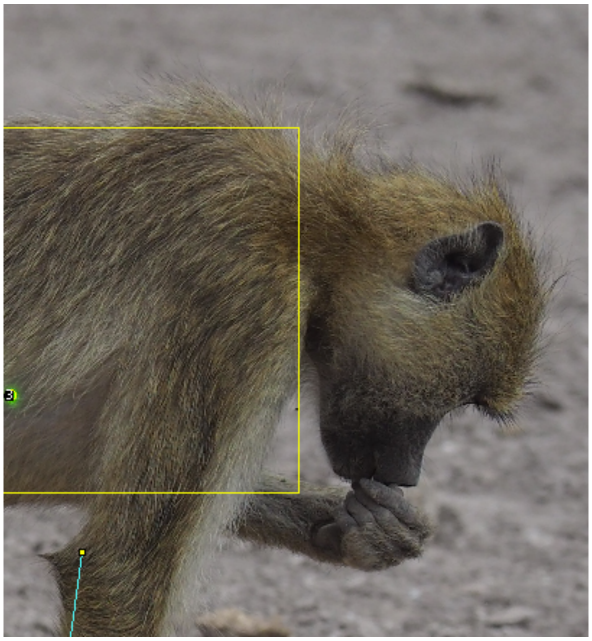
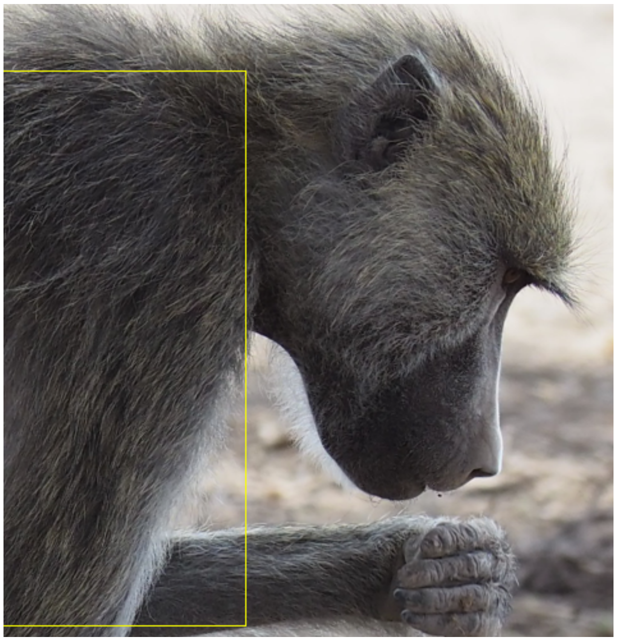
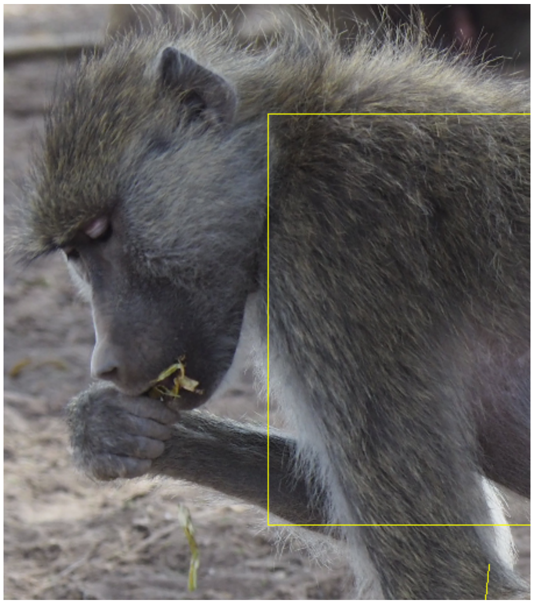

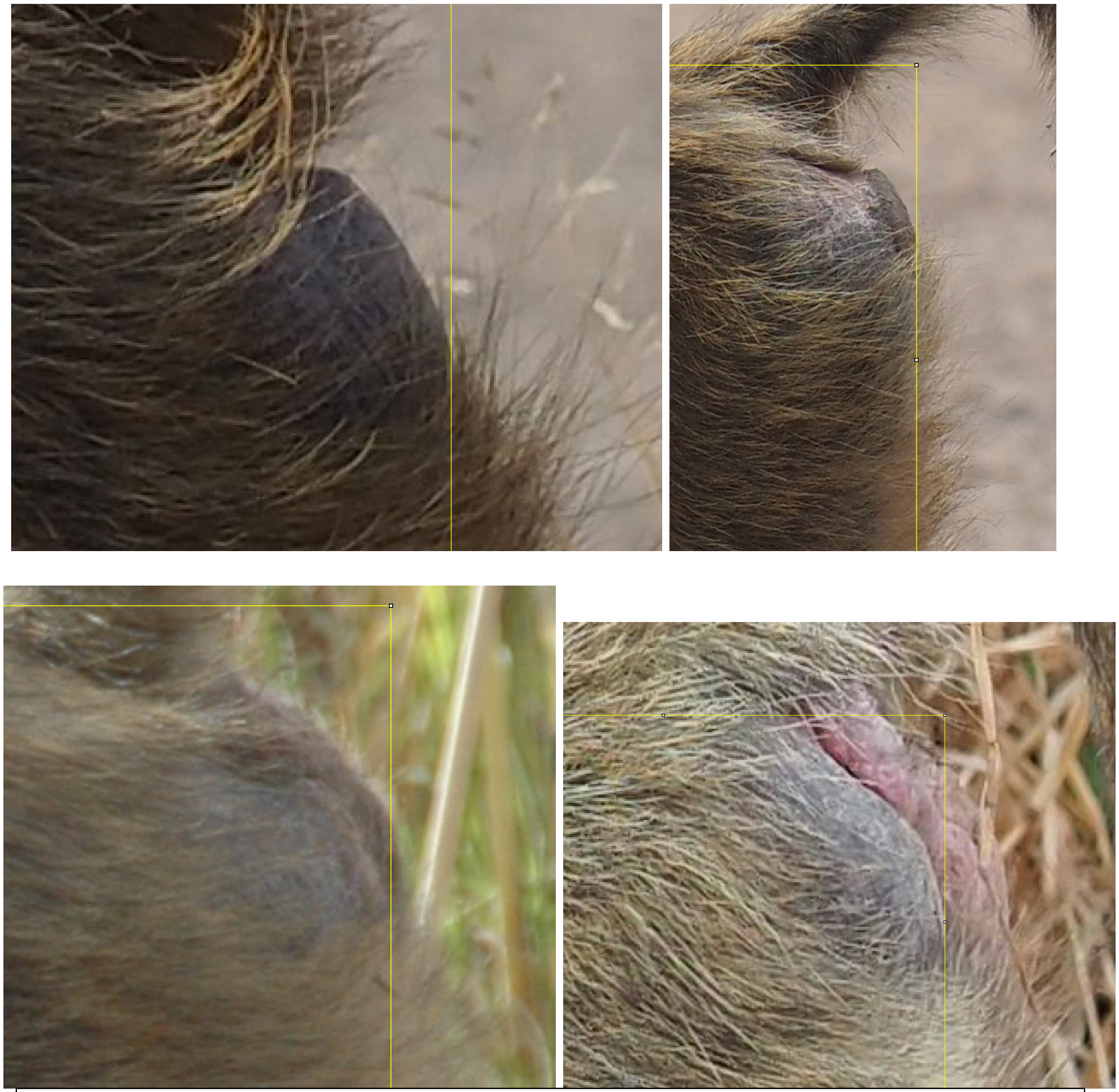

Left: The female has a small sexual swelling between the callosities. Instead of measuring the swelling, we stop the measurement at the end of the callosity.
Right: A much larger swelling is a little easier to identify and measure around.

LOWER LEG

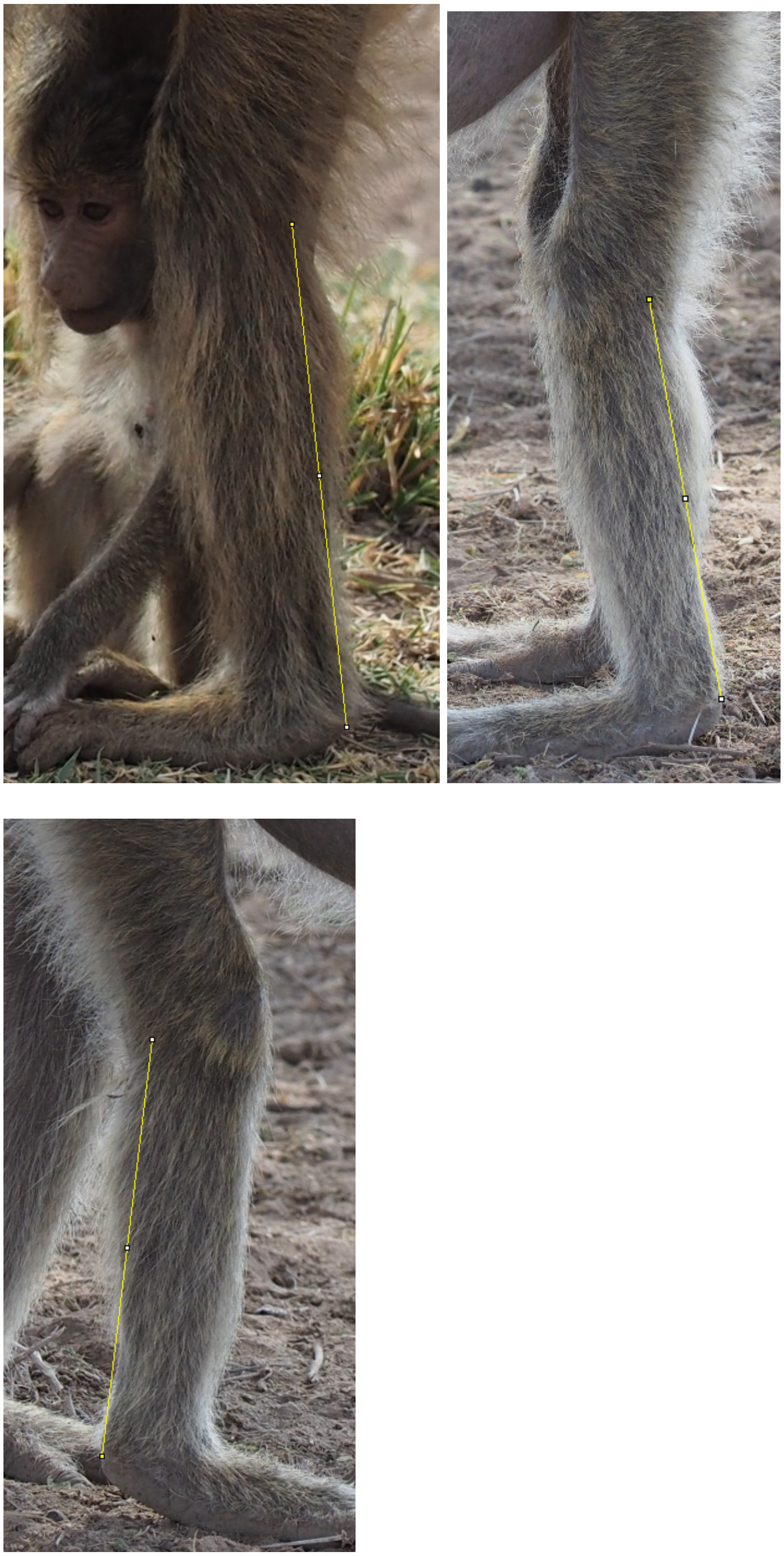

FEMUR

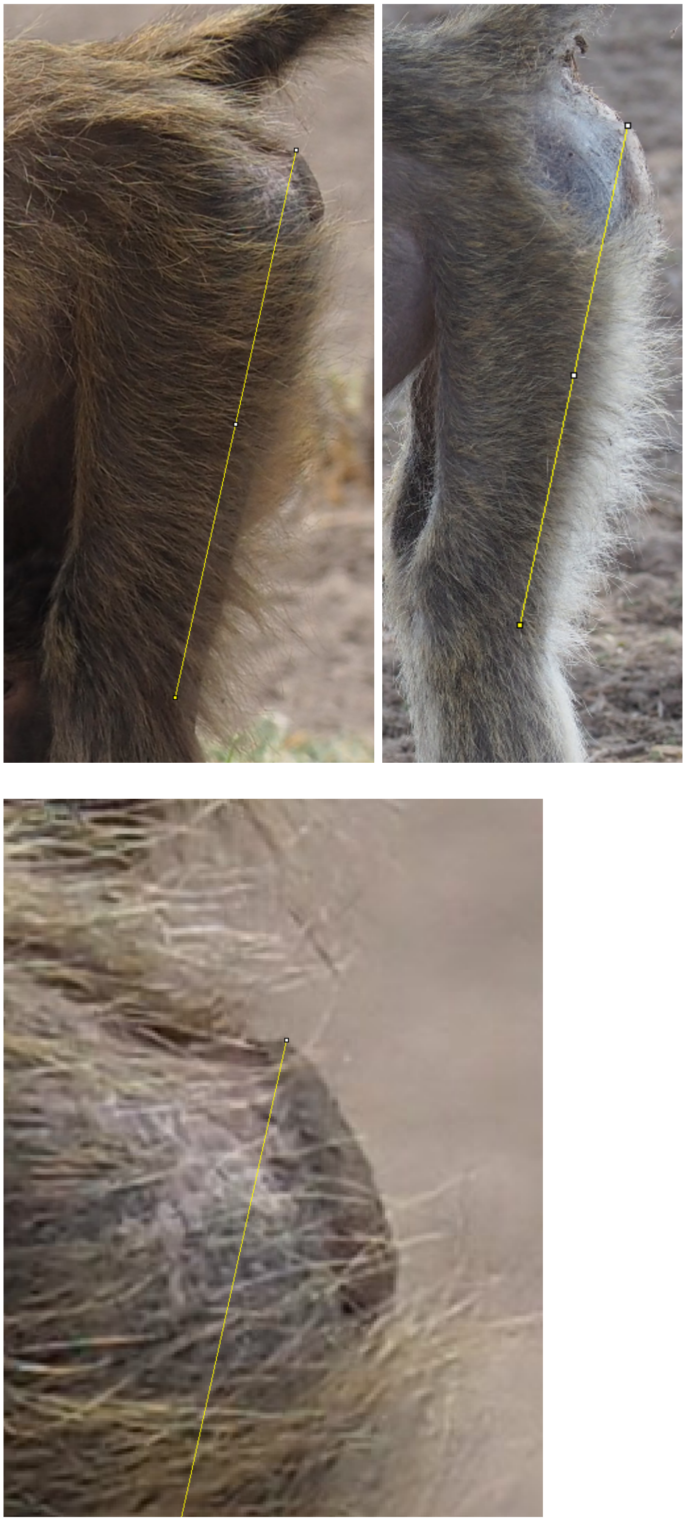

FOREARM

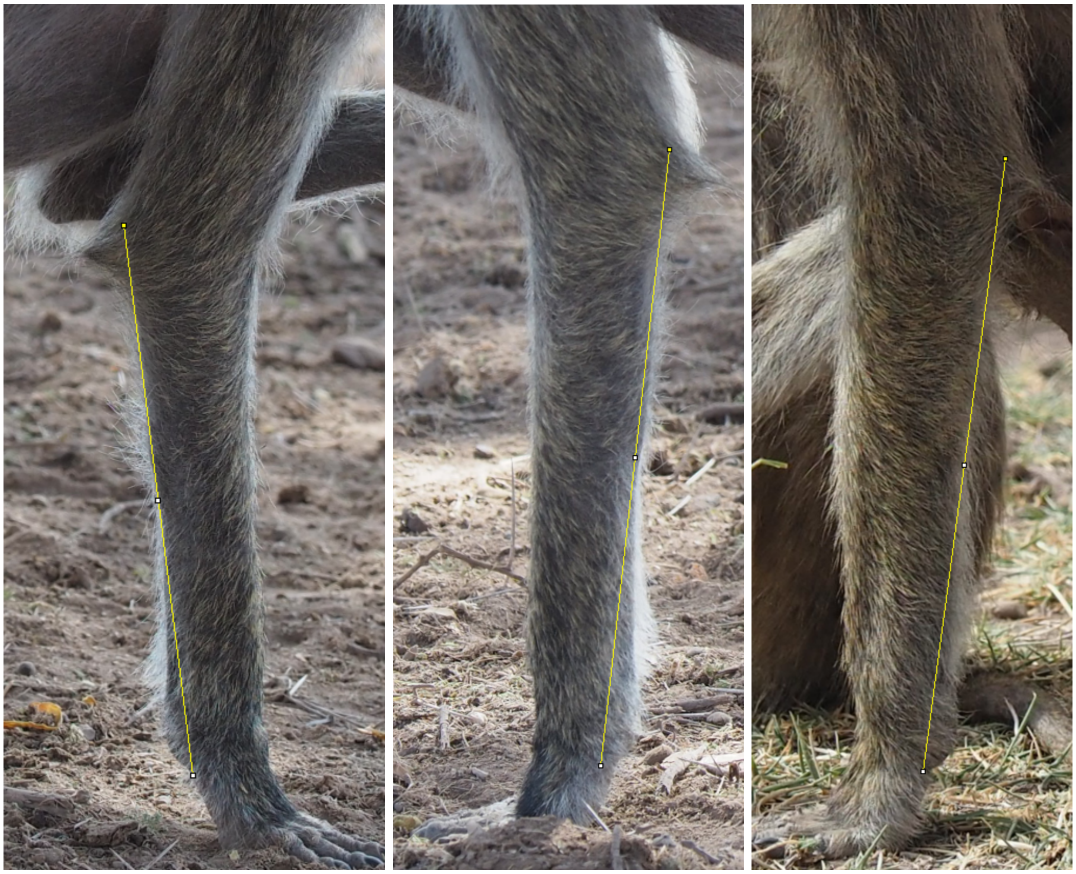
